# Segregated localization of target-SNARE proteins within presynaptic terminals of Munc18-1 deficient photoreceptors

**DOI:** 10.1101/2024.12.19.629437

**Authors:** Mengjia Huang, Chun Hin Chow, Jason Charish, Karen Indrawinata, Hidekiyo Harada, Valerie A. Wallace, Philippe P. Monnier, Shuzo Sugita

## Abstract

Sec1/Munc18 family proteins are essential for SNARE-mediated vesicular exocytosis. However, where SNARE proteins are localized in Munc18-1 deficient presynaptic terminals remains unclear due to the rapid degeneration of neurons lacking Munc18-1. Here, we found that removing Munc18-1 from photoreceptor cells did not result in major cellular loss until postnatal day 14, which allowed us to investigate the role of Munc18-1 in endogenous presynaptic terminals. In the absence of Munc18-1, even before major photoreceptor cell degeneration, functional impairments were present. While Munc18-1 was not required for the pre-synaptic enrichment of the t-SNARE proteins syntaxin-3 and SNAP-25, it played a critical role in their proper localization. In wild-type conditions, t-SNAREs are highly colocalized. However, in the absence of Munc18-1, their distribution becomes strikingly segregated. Immuno-electron microscopy revealed that without Munc18-1, syntaxin-3 is retained within various organelle membranes rather than being targeted to synaptic plasma membranes. These findings provide the first evidence that Munc18-1 is important to prevent segregation of syntaxin-3 and SNAP-25 within presynaptic terminals.

## Introduction

The retina is the light sensitive tissue at the back of the eye and is made up of 5 types of neurons. These cells stratify into three distinct nuclear layers that are separated by two synaptic “plexiform” layers for cell-to-cell communication (Seung and Sümbül, 2014; Sanes and Zipursky, 2010). Visual signals are transduced by photoreceptor cells located in the outer nuclear layer (ONL), passed onto bipolar cells in the inner nuclear layer (INL), and then onto ganglion cells in a three-neuron chain (Kolb, 1995). Retinal synapses are highly ordered, complete their development postnatally, and form well-defined connections, making the retina exceptionally useful for the study of synapse development, refinement, and maintenance.

In the outer retina, photoreceptor cells adjust glutamate release to convey light information to their postsynaptic bipolar cells using special synapses termed ribbon synapses. Synaptic vesicles are tethered to this proteinaceous ribbon structure in preparation for exocytosis. The SNARE complex, composed of isoforms of syntaxin, synaptosomal associated protein (SNAP), and synaptobrevin/vesicle-associated membrane protein (VAMP), is the main machinery executing membrane fusion and exocytosis for neurotransmitter release at conventional synapses (Südhof and Rizo, 2011; Südhof and Rothman, 2009; Südhof, 2004). Despite previous suggestions that exocytosis at ribbon synapses may occur through SNARE-independent mechanisms (Nouvian et al., 2011), it is now evident that exocytosis at photoreceptor and inner hair cell ribbon synapses is SNARE dependent, albeit with alternate SNARE isoforms (Calvet et al., 2022). Specifically, we and others established that syntaxin-3 and SNAP-25 are the obligate target SNARE (t-SNARE) proteins utilized for synaptic transmission as well as outer segment dynamics in photoreceptor cells (Kakakhel et al., 2020; Janecke et al., 2021; Huang et al., 2024). However, how SNARE-mediated exocytosis is regulated at photoreceptor cell ribbon synapses remains unknown.

Complementary to SNARE proteins, Sec1/Munc18-like (SM) proteins help to orchestrate membrane fusion in conventional synapses. Munc18-1, also known as STXBP1, was originally described as a syntaxin-1 binding protein (Hata et al., 1993). Munc18-1 is required for neuron survival and neurotransmitter release as shown by Munc18-1 null mice (Verhage et al., 2000) and Munc18-1 null neuron culture (Vardar et al., 2016; Deák et al., 2009). However, because Munc18-1 deficiency compromises neuron survival, how it organizes SNARE proteins in presynaptic nerve terminals is less understood. For example, Munc18-1 appears to be required for syntaxin-1 expression in neurons, where in the absence of Munc18-1, syntaxin-1 levels are reduced by 70%, although the remaining syntaxin-1 is reported to be correctly targeted to synapses (Verhage et al., 2000). On the other hand, a perturbation in plasmalemmal syntaxin-1 has been demonstrated in Munc18-1 knockdown or Munc18-1/-2 double knockdown PC12 cells (Han et al., 2009; Arunachalam et al., 2008). Similar changes in syntaxin-1 distribution have also been reported in soma of neurons that are differentiated from patient-derived Munc18-1 heterozygous mutant iPSCs (Yamashita et al., 2016). The rapid death of neurons makes investigating Munc18 function and protein interactions in primary neurons challenging, but surrogate systems such as PC12 and iPSCs lack true synapses. Thus, how Munc18-1 regulates syntaxin transport to presynaptic nerve terminals remains elusive. Further, the consequences of disrupting Munc18-1 and syntaxin interaction on other SNARE proteins or SNARE complex localization are unclear.

A heterozygous mutation (H16R) in Munc18-1 is known to be associated with congenital nystagmus in humans (Li et al., 2020). Nevertheless, the exact roles and functions of Munc18-1 in photoreceptor cells and vision remain unknown. Synaptic vesicle exocytosis must be carefully and precisely regulated in photoreceptors, as neurotransmitter release is tonic rather than phasic. Light stimulation hyperpolarizes photoreceptor cells, leading to reduced glutamate release. Sustaining the massive glutamate release in photoreceptor cells under dark conditions, along with mediating rapid modulation of vesicle release in response to light relies on the activity of key regulatory proteins. Therefore, it is crucial to identify Munc18-1’s role in SNARE protein targeting in unconventional photoreceptor ribbon synapses.

To address these questions, we generated photoreceptor-specific Munc18-1 conditional knockout (cKO) mice, wherein CRX-driven cre-recombinase induces the removal of Munc18-1 from photoreceptors and a subset of bipolar cells starting on E12.5. Then using behavioral and functional vision tests, combined with histology and ultrastructural analyses, we examine the consequences of Munc18-1 loss in photoreceptor cells. Despite significant impairment to photoreceptor cell functioning by postnatal day 14 (P14), remarkably, the majority Munc18-1 cKO photoreceptors survived until P14, providing an opportunity to investigate the interplay between Munc18-1 and other SNARE proteins in vivo. Here, we address three key questions: i) whether SNARE-dependent ribbon synapse exocytosis is mediated by Munc18-1, ii) whether syntaxin-3 can be targeted to synapses in the absence of Munc18-1, and iii) whether syntaxin-3 can localize with SNAP-25 following the removal of Munc18-1. Our results demonstrate that Munc18-1 is essential for photoreceptor cell function and survival and that Munc18-1 function is required to prevent the segregation of syntaxin and SNAP-25 on the plasma membrane.

## Results

### Munc18-1 removal from photoreceptors leads to retina thinning and loss of visual function

To investigate the roles of Munc18-1 in photoreceptor neurons, we specifically removed Munc18-1 from photoreceptor cells and a subset of bipolar neurons by crossing Munc18-1 flox mice (Park et al., 2017) with CRX-cre mice (Prasov and Glaser, 2012). Because CRX-Cre primarily targets photoreceptor cells, we refer to this mouse line as a photoreceptor-specific Munc18-1 knockout. We started with *in vivo* retinal imaging and behavioral tests to identify gross morphological and functional changes at P22 when photoreceptor cell maturation is complete. Optical coherence tomography (OCT) imaging showed that Munc18-1 cKO mice had reduced retinal thickness by P22 (Supplemental Fig. 1A). Electroretinogram (ERGs) analysis under dark-adapted (scotopic) and light-adapted (photopic) conditions showed that Munc18-1 cKO photoreceptor cells did not hyperpolarize in response to light (no A-wave), and also failed to induce a downstream B-wave (Supplemental Fig. 1B). Finally, adult Munc18-1 cKO mice had a profound reduction in visual acuity, as measured by the optokinetic tracking response when compared to the controls (Supplemental Fig. 1C). Therefore, we next moved to section histology to analyze the eyes of Munc18-1 cKO and littermate control mice to corroborate the changes we saw by OCT. We first check Munc18-1 control and cKO mice at the same timepoint when electroretinograms and OCT scans were performed – histological analysis of the retina of Munc18-1 cKO mice at P22 revealed a significant reduction in retina thickness, confirming the OCT results. We found a severe loss in total retina thickness in Munc18-1 mice, as virtually no photoreceptor cells remained at P22 (Supplemental Fig. 1D), confirming the retina thickness decrease we observed by optical coherence tomography (Supplemental Fig. 1A).

### Munc18-1 deficient photoreceptors degenerate between P14 and P22

To understand why photoreceptor cells do not survive to P22 in Munc18-1 cKO mice, we examined photoreceptor cells at younger timepoints (P9, P12, and P14) to further investigate structural and functional changes. We first confirmed the successful removal of Munc18-1 using a polyclonal Munc18-1 antibody (Fig. 1A). In control sections, Munc18-1 was abundantly present in photoreceptor cells and their synaptic terminals, as Munc18-1 expression was seen in the outer nuclear layer (ONL) and colocalized with PSD95 which identifies the presynaptic region of the outer plexiform layer (OPL). Munc18-1 was found throughout the outer plexiform layer, and also strongly observed throughout the inner nuclear layer (INL), inner plexiform layer (IPL), and ganglion cell layer (GCL)/retinal nerve fiber layer. However, in Munc18-1 cKO mice, Munc18-1 fluorescence was largely absent from the photoreceptors of the ONL and the Munc18-1 signal colocalizing with presynaptic marker PSD95 was mostly absent, confirming effective Munc18-1 knockout in photoreceptor cells (Fig. 1A). Munc18-1 was also absent in the apical INL, indicating Munc18-1 removal from a subset of bipolar cells (Kakakhel et al., 2020). However, Munc18-1 remained abundantly expressed throughout the basal INL and IPL. We were able to confirm this independently using a second monoclonal Munc18-1 antibody and similarly observed a significant decrease in Munc18-1 signal fluorescence in photoreceptor cells and bipolar cells of the knockout sections (Fig. 1B), while basal INL and IPL Munc18-1 remained intact. Examining sections at P12 and P14, we found similar photoreceptor and bipolar cell removal of Munc18-1 at both ages (Fig. 1C). Despite Munc18-1 being efficiently removed in P9 photoreceptor cells, we did not observe a decrease in photoreceptor layer thickness at P9 in Munc18-1 cKO mice when compared to age-matched controls (Fig. 1D) and we did not observe significant photoreceptor loss leading up to P14, indicating photoreceptor loss largely occurs between P14 and P22 in Munc18-1 cKO retinas.

**Figure 1.**
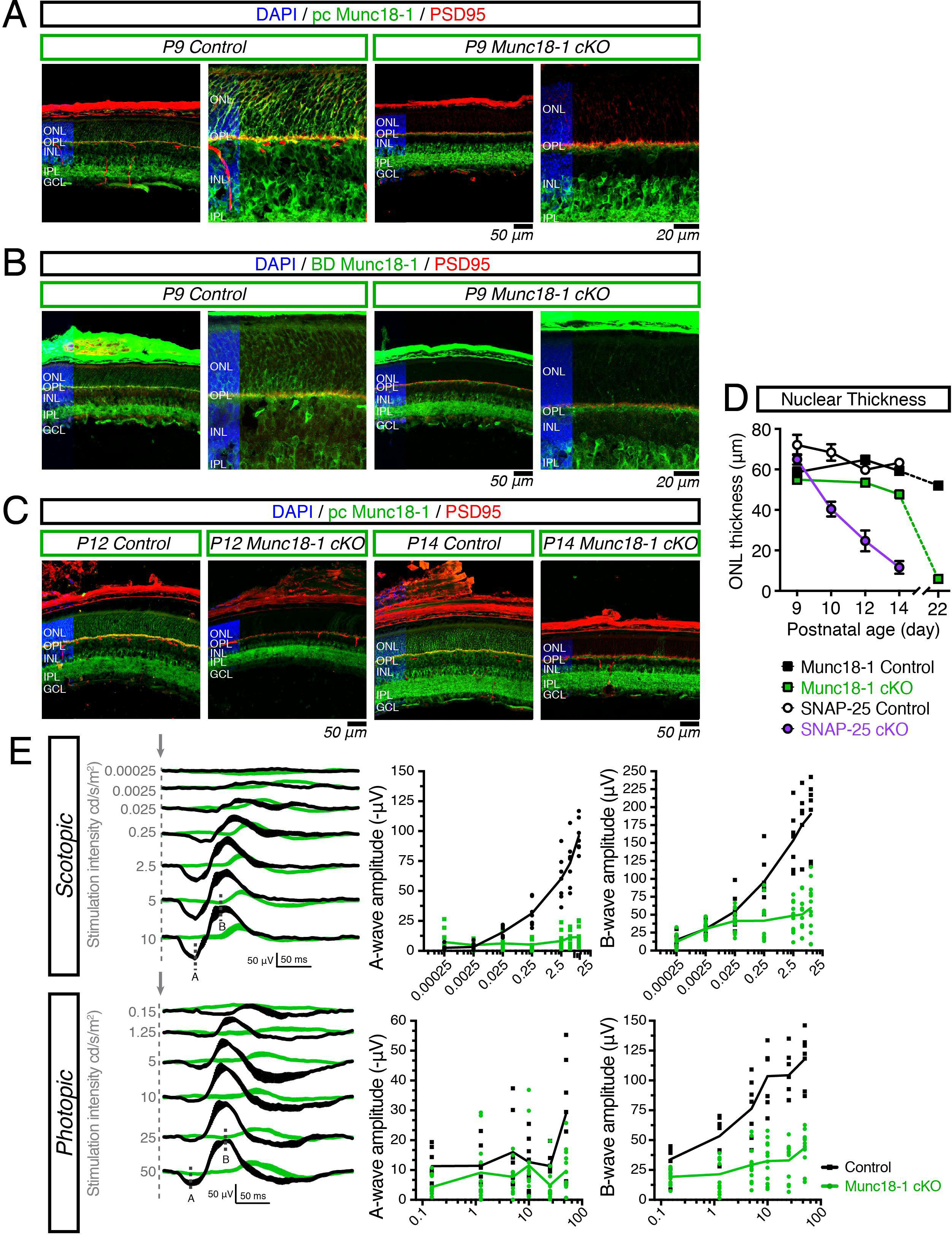
Effective removal of Munc18-1 by CRX-cre leads to degeneration between postnatal day 14 and 22. (A) Antibody staining for Munc18-1 (green) using rabbit polyclonal antibody against PSD95 (red) for presynaptic terminals in P9 control and P9 Munc18-1 retina. Munc18-1 is present throughout the retina in the outer nuclear layer (ONL), outer plexiform layer (OPL), inner nuclear layer (INL), inner plexiform layer (IPL), and ganglion cell layer (GCL). Munc18-1 fluorescence is lost in the photoreceptor and bipolar cells of Munc18-1 cKO mice. Retina region for all represented images is 0.5–1 mm away from optic nerve. Scale bar = 50 μm and scale bar = 20 μm. (B) Antibody staining for Munc18-1 (green) using mouse monoclonal antibody and PSD95 (red) in P9 control and Munc18-1 cKO retina. A decrease in Munc18-1 fluorescence is signal is observed in Munc18-1 cKO mice. Retina region for all represented images is 0.5–1 mm away from optic nerve. Scale bar = 50 μm and scale bar = 20 μm. (C) Antibody staining for Munc18-1 (green) using rabbit polyclonal antibody against PSD95 (red) for presynaptic terminals in P12 and P14 control and Munc18-1 cKO retina. (D) Quantification of outer nuclear layer (ONL) thickness in control, Munc18-1 cKO (green, squares) mice and SNAP-25 cKO (purple, circles) mice. SNAP-25 cKO photoreceptors are rapidly lost between P9 and P14 while Munc18-1 cKO photoreceptors are rapidly lost between P14 and P22. (E) Averaged scotopic (dark-adapted) and photopic (light-adapted) electroretinogram traces across 7 light intensities in P14 control and P14 Munc18-1 cKO mice. Arrow indicate stimulation onset, A indicates a-wave, B indicates b-wave. Munc18-1 cKO mice exhibit diminished light responses at all light intensities. One eye was excluded from a control mouse due to eyelid deformity.

While Munc18-1 cKO ultimately leads to photoreceptor cell degeneration, photoreceptors were largely maintained at P14, which is the earliest possible time to perform electroretinograms. We took advantage of this and tested the ERG response of Munc18-1 cKO mice at P14 at eye opening (Fig. 1E). We found that despite the majority of photoreceptor cells being maintained in Munc18-1 cKO mice, their ERG response at P14 is still largely reduced (Fig. 1E). Our results suggest that Munc18-1 is necessary for light responsiveness and synaptic communication of photoreceptors.

We previously generated SNAP-25 conditional knockout mice, using the same CRX-driven cre-recombinase; and SNAP-25 cKO photoreceptor cells have rapid degeneration that is complete by P14 (Huang et al., 2024). This is in contrast to the Munc18-1 cKO mice, where photoreceptor cells are largely maintained until P14. Using electron micrographs (EM), we found in SNAP-25 cKO mice that the photoreceptor degeneration appeared to start from the lowest strata of the outer nuclear layer (Huang et al., 2024). As SNAP-25 cKO and Munc18-1 cKO exhibited a different timescale of neurodegeneration, we further look into the spatial pattern of cell death. We compared DNA fragmentation using TUNEL in SNAP-25 cKO and Munc18-1 cKO mice. Similar to what we observed by EM, we saw TUNEL staining for DNA fragmentation beginning in the lowest strata of the ONL in SNAP-25 cKO mice (Supplemental Fig. 2), and when we divided the outer nuclear layer into thirds, TUNEL-positive cells were significantly increased in the lowest third of the ONL in SNAP-25 cKO mice. In contrast, Munc18-1 cKO TUNEL-positive cells were evenly distributed in the outer nuclear layer (Supplemental Fig. 2). The differential distribution of TUNEL signal and different time courses of degeneration suggest that photoreceptor cell degeneration in Munc18-1 cKO mice and SNAP-25 cKO mice may occur through different mechanisms.

### Altered syntaxin-3 localization in the soma and outer segment region, but not the synapse, of Munc18-1 deficient photoreceptor cells

It has previously been reported that in the absence or mutation of Munc18-1, syntaxin-1 is mislocalized in the soma of PC12 cells and iPSC-derived neurons (Han et al., 2009; Yamashita et al., 2016; Arunachalam et al., 2008). Therefore, we investigated the localization of syntaxin-3 in Munc18-1 cKO retinas. In the control retina, Munc18-1 and syntaxin-3 were both found throughout photoreceptor cells in their outer segments, synapses, and with a plasmalemmal pattern surrounding photoreceptor soma. As a result, a strong colocalization of Munc18-1 and syntaxin-3 was present in the control sections (Fig. 2A). However, in P12 Munc18-1 cKO retina sections, plasmalemmal syntaxin-3 was strongly decreased except for small areas where cre-recombination was less effective, and Munc18-1 remained present (Fig. 2A).

**Figure 2.**
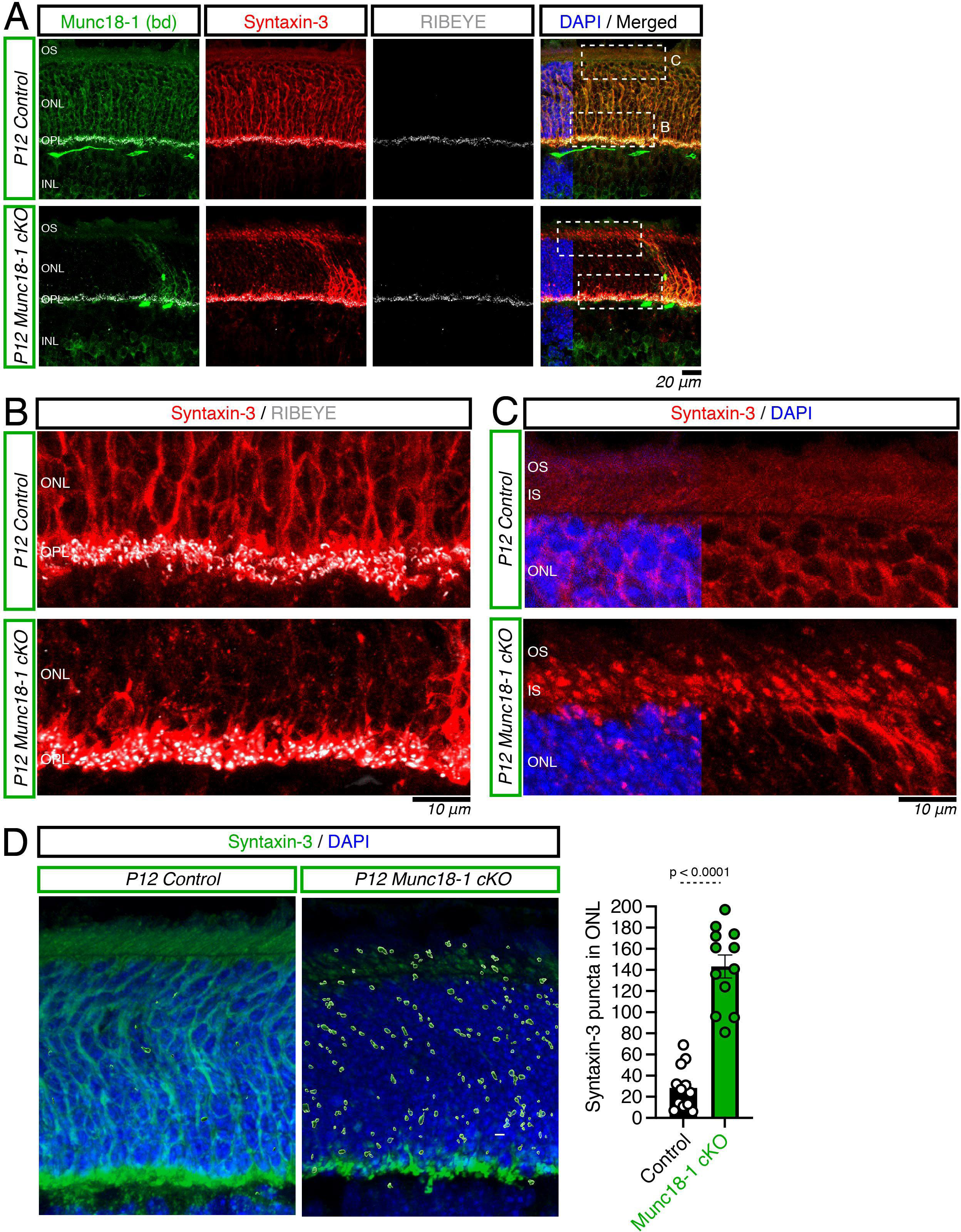
Syntaxin-3 mislocalisation in Munc18-1 cKO mice. (A) Munc18-1 (green), syntaxin-3 (red), and RIBEYE (white) stain in P12 control and Munc18-1 cKO retina. Munc18-1 and syntaxin-3 show strong colocalization in control retina and is found in photoreceptor outer segments, synapses, and in nuclear plasmalemmal. in Munc18-1 cKO retinas, regions where Munc18-1 is absence also have decreased plasmalemmal syntaxin-3 signal. Syntaxin-3 remains abundantly present in Munc18-1 cKO synapses. (B) Magnified synaptic region from (A), in control retina, syntaxin-3 is present surrounding photoreceptor nuclei and in the synapses. In Munc18-1 cKO synapses, however plasmalemmal syntaxin-3 signal decreases despite the synaptic syntaxin-3 remaining intact. (C) Magnified outer segment region from (A). Syntaxin-3 is uniformly found through the ciliary outer segment structures of control photoreceptors. In Munc18-1 cKO retina, the ciliary staining pattern is lost – rather, syntaxin-3 signal appears punctate and preferentially localized in the inner segment (IS) region. Syntaxin-3 punctate signal is also observed throughout the outer nuclear layer. (D) Volume projection of syntaxin-3 (green) in P12 control and Munc18-1 cKO mice. A decrease in continuous syntaxin-3 plasmalemmal localization is present in Munc18-1 cKO mice. Trapped syntaxin-3, identified as bright puncta, are increased in Munc18-1 cKO retina when compared with controls, t_(22)_ = 9.272; p < 0.0001.

Concurrent with the disappearance of syntaxin-3 from the plasma membrane, we observed accumulation of syntaxin-3 in the perinuclear and outer segment regions of Munc18-1 cKO photoreceptors (Fig. 2A, 3C). That is, bright puncta of syntaxin-3 was observed within the outer nuclear layer and in the outer segment region of Munc18-1 deficient photoreceptor cells (Fig. 2C). Using 3D reconstruction, we were able to visualize and quantify the perinuclear syntaxin-3 accumulations in Munc18-1 cKO retina sections (Fig. 2D) where we saw a significant increase in syntaxin-3 puncta in the ONL. Therefore, without Munc18-1, syntaxin-3 at the photoreceptor cell soma was not properly localized.

Despite the decrease of plasmalemmal syntaxin-3, syntaxin-3 still localized with RIBEYE, a marker for the synaptic ribbon, in Munc18-1 cKO synaptic terminals (Fig. 2A-B). Thus, our results suggest that synaptic transport of syntaxin-3 to the photoreceptor synapse is preserved in the absence of Munc18-1. This result is consistent with the previous findings from embryonic day 18 (E18) brain slices of Munc18-1 null mice, where the remaining syntaxin-1 was targeted to synaptic regions (Verhage et al., 2000).

Changes in syntaxin-3 localization in the photoreceptor cells, particularly accumulations in the outer segments, could be an artefact of severe degeneration (Baker et al., 2008). Therefore, we examined syntaxin-3 localization in SNAP-25 cKO mice, which also exhibit severe photoreceptor cell degeneration (Huang et al., 2024). Both t-SNARES syntaxin-3 and SNAP-25 colocalized in control sections as expected in control sections (Supplemental Fig. 3). In SNAP-25 cKO mice, syntaxin-3 continued to be present in the outer segments, surrounding photoreceptor nuclei plasmalemma, and in the synapses (Supplemental Fig. 3). Therefore, we conclude that the changes in syntaxin-3 staining observed in Munc18-1 cKO retinas are not the consequence of severe degeneration, and rather a specific effect of Munc18-1 deletion.

### Removal of Munc18-1 leads to dissociation in the location of t-SNARE proteins within presynaptic terminals

Since the localization of syntaxin-3 is changed in Munc18-1 deficient photoreceptors, we next asked whether this change is specific to syntaxin-3, or if Munc18-1 removal affected the localization of the other t-SNARE, SNAP-25. Target-SNARE partners syntaxin-3 and SNAP-25 were both abundantly present in photoreceptor cells and showed strong colocalization in the control retina as expected. We observed no differences in SNAP-25 expression pattern between the Munc18-1 cKO retina versus control retina. SNAP-25 was abundantly found in the outer segments, surrounding photoreceptor nuclei, in the outer plexiform layer, and throughout the inner retina (Fig. 3A). We conclude that SNAP-25 localization in photoreceptors is unaffected by Munc18-1 deletion.

**Figure 3.**
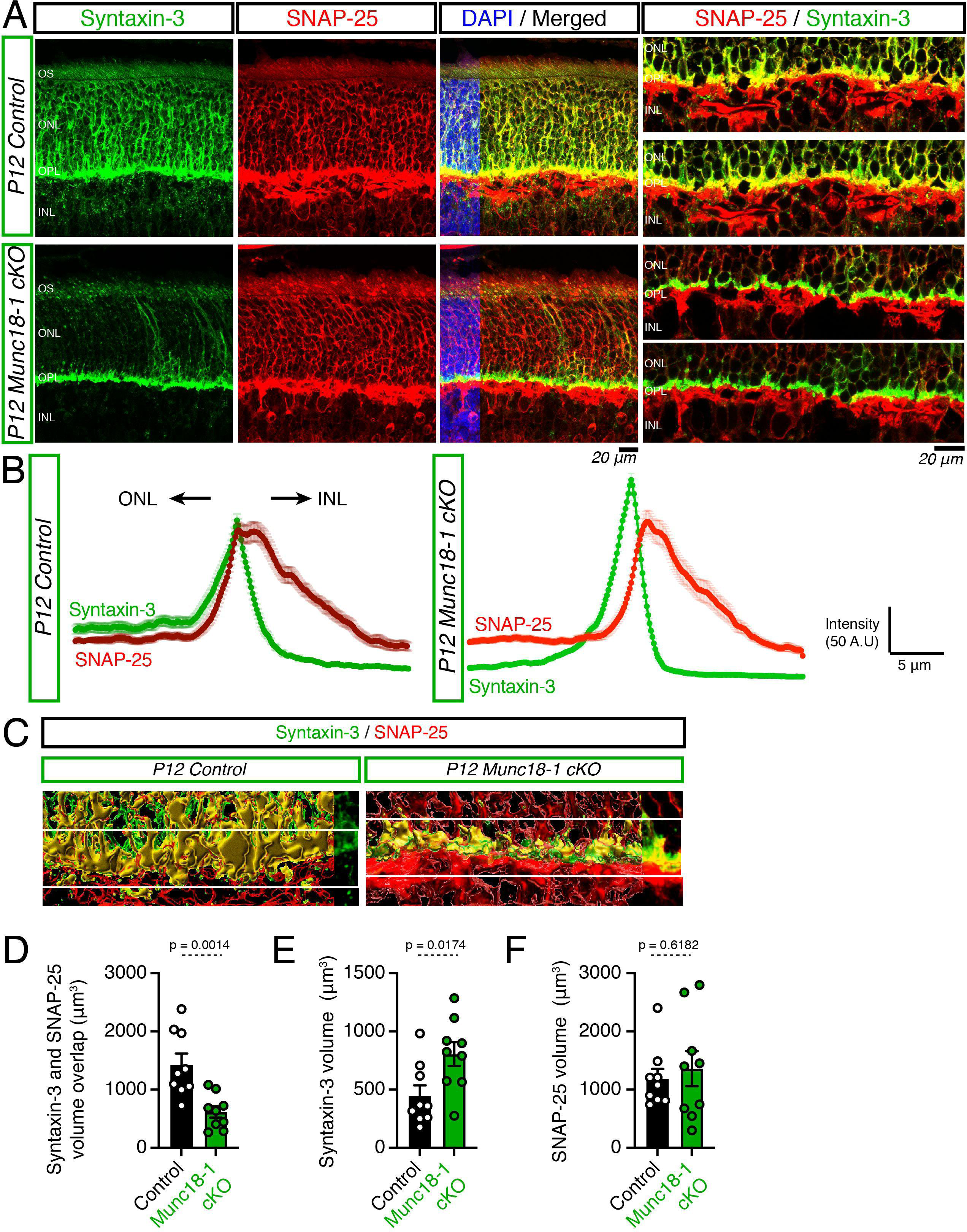
Dissociation between t-SNAREs Syntaxin-3 and SNAP-25 in Munc18-1 cKO. (A) Syntaxin-3 (green) and SNAP-25 (red) immunostaining in P12 control and Munc18-1 cKO retinas. Syntaxin-3 and Munc18-1 strongly colocalize in control retina sections (yellow). Syntaxin-3 loses somatic locale in Munc18-cKO retinas while SNAP-25 signal remains unchanged in Munc18-1 cKO retina. Synaptic Syntaxin-3 and SNAP-25 no longer colocalize together in Munc18-1 cKO retinas, clearly visualized in magnifications of single z-planes presented on right. (B) Intensity profile of syntaxin-3 (green) and SNAP-25 (red) in P12 control and Munc18-1 cKO retinas. An overlap in syntaxin-3 and SNAP-25 intensity is observed in control retinas, consistent with the observed colocalization. However, the peaks of syntaxin-3 and SNAP-25 signal intensity are shifted and no longer overlap in Munc18-1 cKO retinas and is consistent with the loss of outer plexiform layer colocalization observed in (A). (C) Representative 3D volume projections of the outer plexiform layer in P12 control and Munc18-1 cKO retina. A decrease in the volume of syntaxin-3 and SNAP-25 overlap is observe in Munc18-1 cKO retina. (D) Quantifications of syntaxin-3 and SNAP-25 volume overlap, t_(16)_ = 3.862; p = 0.0014. (E) Quantifications of syntaxin-3 volume not colocalized with SNAP-25, t_(16)_ = 2.652; p = 0.0174. (F) Quantifications of SNAP-25 volume not colocalized with syntaxin-3, t_(16)_ = 0.508; p = 0.6182.

Unexpectedly, when we looked at syntaxin-3 and SNAP-25 localization in Munc18-1 deficient retinas, we noted a striking difference – synaptically targeted syntaxin-3 did not colocalize with SNAP-25 in the outer plexiform layer, rather, syntaxin-3 fluorescence appeared to be apical to the SNAP-25 signal (Fig. 3A). Using ImageJ, we generated intensity profiles for both syntaxin-3 and SNAP-25 in control and Munc18-1 cKO sections to quantitatively analyze the immunofluorescence signal (Fig. 3B). In control retina, the intensity profile of syntaxin-3 largely overlapped with the SNAP-25 signal and the peak of syntaxin-3 intensity was within the peak of SNAP-25 fluorescence. However, an apical shift can be observed in the peak of syntaxin-3 signal in Munc18-1 deficient retinal regions, wherein the syntaxin-3 peak shifts towards to ONL relative to the peak of SNAP-25 (Fig. 3B). We additionally generated volume renderings of control and Munc18-1 cKO sections and confirm a significant decrease in the volume overlap between syntaxin-3 and SNAP-25 (Fig. 3C-D). Furthermore, the proportion of syntaxin-3 that did not colocalize with SNAP-25 in Munc18-1 cKO was significantly increased in Munc18-1 deficient retinas (Fig. 3E), supporting the dissociation of these two t-SNARE proteins. The volume of SNAP-25 signal alone was not significantly different between control or Munc18-1 cKO retina (Fig. 3F). Therefore, while the synaptic transport of syntaxin-3 remained intact in Munc18-1 deficient photoreceptor cells, the decreased colocalization between syntaxin-3 and its cognate t-SNARE partner suggests that in the absence of Munc18-1, there are alterations in t-SNARE protein localization.

### Changes in Munc18-1 SNARE localization are specific to syntaxin-3

Two possibilities exist for the loss of colocalization between syntaxin-3 and SNAP-25: first, there are changes in the presynaptic localization of SNAP-25; second, despite syntaxin-3 being targeted to the synaptic region, syntaxin-3 is not reaching the synaptic membrane where the photoreceptor active zone is. To test the first possibility, we first co-stained SNAP-25 with the ribbon marker RIBEYE. SNAP-25 was found in the OPL and colocalized with RIBEYE in Munc18-1 cKO photoreceptor terminals (Fig. 4A), excluding the possibility that SNAP-25 presynaptic localization is altered. Next, we co-stained syntaxin-3 with RIBEYE to investigate the localization of syntaxin-3. In control sections, syntaxin-3 fluorescence was found throughout the outer plexiform layer and extended past the RIBEYE signal, however, in Munc18-1 cKO sections, syntaxin-3 was restricted to the presynapse and the signal did not extend beyond the RIBEYE signal (Fig. 4B). Therefore, while SNAP-25 localization appears unchanged in P12 Munc18-1 cKO retina, syntaxin-3 localization is disrupted in the absence of Munc18-1.

**Figure 4.**
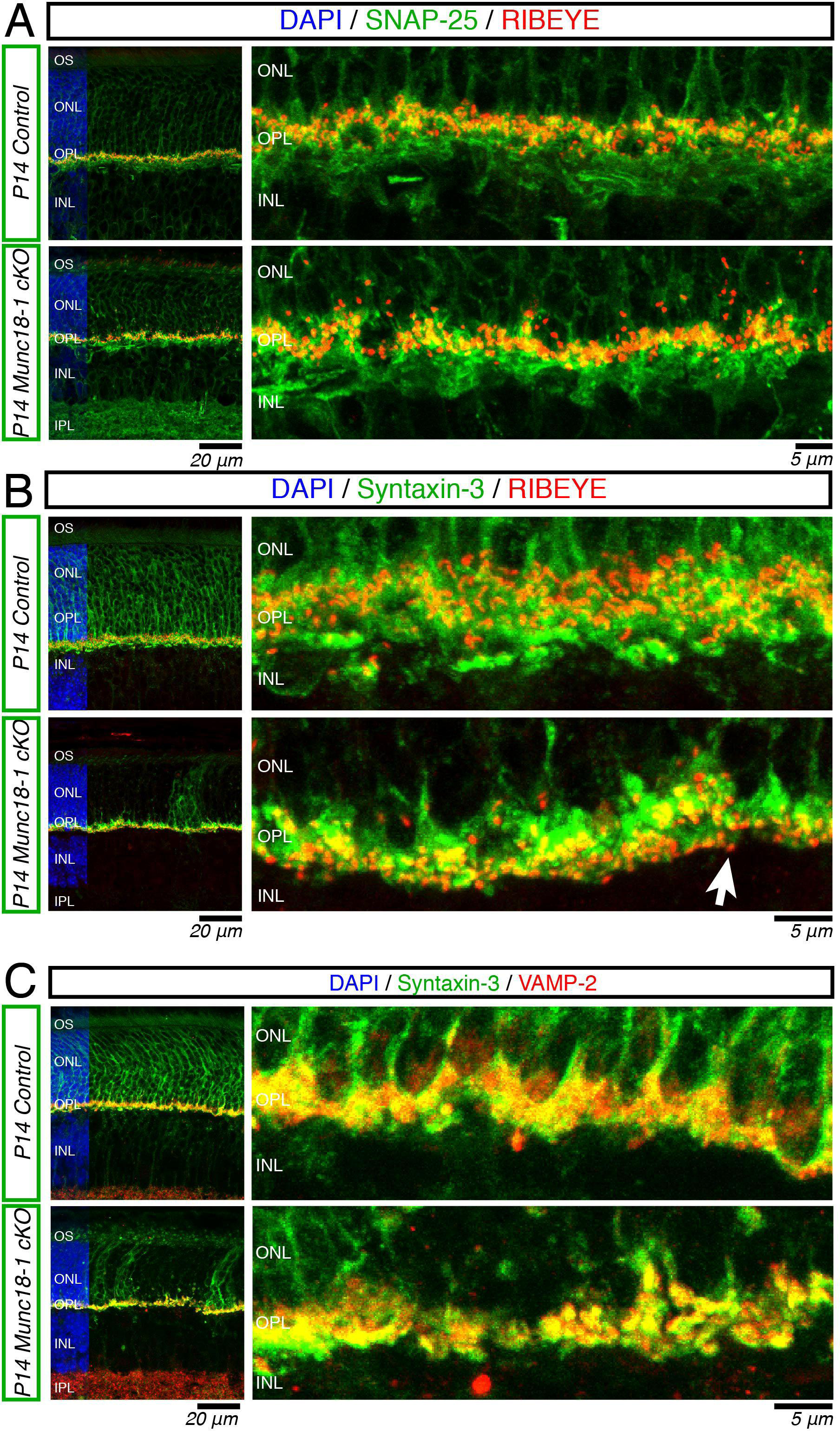
Changes in SNARE mislocalisation in Munc18-1 cKO is specific to syntaxin-3. (A) Confirmation that SNAP-25 localization is unchanged in Munc18-1 cKO. SNAP-25 (green) remains strongly colocalized with presynaptic RIBEYE (red) in Munc18-1 cKO retina. (B) Syntaxin-3 synaptic mislocalisation in Munc18-1 cKO. Syntaxin-3 (green) staining pattern is altered in P14 Munc18-1 cKO retinas – syntaxin-3 signal no longer extends past the presynaptic RIBEYE signal in Munc18-1 cKO retinas. (C) Syntaxin-3 (green) and synaptobrevin-2 (red) in control and Munc18-1 cKO retinas. Synaptic syntaxin-3 colocalizes with synaptobrevin-2 in both control and Munc18-1 retinas, although diffuse somatic syntaxin-3 is lost in Munc18-1 cKO retinas. Syntaxin-3 signal is largely in bright spherical puncta.

After investigating the two target SNAREs, syntaxin-3 and SNAP-25, we next looked at the vesicular SNARE protein synaptobrevin-2 to determine whether the mislocalisation was specific to syntaxin-3 or also included synaptobrevin-2. We observed the presynaptic presence of synaptobrevin-2 in P9 control and Munc18-1 cKO sections – synaptobrevin-2 colocalized with presynaptic PSD95 in both control and Munc18-1 cKO photoreceptors suggesting that the presynaptic localization of synaptobrevin-2 was unchanged by Munc18-1 removal (Supplemental Fig. 4A). Next, we co-stained syntaxin-3 with synaptobrevin-2 to compare their locations – the apical shift and spherical pattern of syntaxin-3 fluorescence signal could suggest that syntaxin-3 is colocalizing/trapped with v-SNARES rather than t-SNARES. In control sections, syntaxin-3 and synaptobrevin colocalized in the presynaptic OPL and we did not observe a shift in syntaxin-3 localization relative to synaptobrevin-2 in Munc18-1 cKO retinas (Fig. 4C). We conclude that Munc18-1 is not required for the synaptic targeting of synaptobrevin-2. Therefore, loss of Munc18-1 specifically affects syntaxin-3, as neither SNAP-25 nor synaptobrevin-2 localization are affected.

### Syntaxin-3 is mislocalized in organelle membranes in Munc18-1 cKO photoreceptors

The segregation of syntaxin-3 and SNAP-25 at the synapse in Munc18-1 cKO suggests that in the absence of Munc18-1 there is a potential subcellular mislocalisation of syntaxin-3. We next performed pre-embedding immuno-electron microscopy for syntaxin-3 on P12 control and Munc18-1 cKO retinas. In control sections, syntaxin-3 signal was observed adjacent to synaptic ribbons and along the synaptic boundary, emphasizing the plasma membrane that composes developing rod spherule synapses (Fig. 5A, B). In Munc18-1 cKO sections, syntaxin-3 staining was absent from the plasma membrane adjacent to synaptic ribbons. When looking for plasma membranes that were adjacent to synaptic ribbons that had strong syntaxin-3 immunogold labelling, we observed 57/67 (85%) strongly tagged membranes in control retina, while this number decreased to 12/68 (18%) in Munc18-1 cKO retina. Rather, syntaxin-3 immunogold labelling was observed in non-synaptic structures, including the membranes of lipidic vacuoles (Fig. 5A), in vacuolated structures (Fig. 5B), and in the endoplasmic reticulum and Golgi appearing structures in Munc18-1 deficient photoreceptor cells (Fig. 5B-C). Notably, synaptic vesicles appeared larger and had additional definition in Munc18-1 deficient photoreceptor terminals, suggesting syntaxin-3 may indeed be trapped with v-SNARES rather than reaching the plasma membrane where its t-SNARE partner resides. A solution of uranyl acetate and lead citrate was applied to the ultrathin EM sections prior to imaging, adding additional contrast to lipids, proteins, membranes, and cell components to help visualize the cellular morphology. We also imaged sections that lacked the contrasting solution. Here, cellular membranes were less apparent, however in control sections, there were membranes near synaptic ribbons that were easily discernable as a result of immunogold labelling (Fig. 5A, green box. This membrane tagging was largely absent in Munc18-1 synapses and plasma membranes near synaptic ribbons were not apparent (Supplemental Fig. 5B, blue boxes). However, there was also an instance where in the Munc18-1 cKO sample where strong syntaxin-3 immunogold labelling was observed synaptically (Supplemental Fig. 5A, green box), likely identifying a synapse where recombination was less effective, emphasizing the lack of synaptic syntaxin-3 labelling in the other four synapses present. In addition to the absence of syntaxin-3 immunolabelling at photoreceptor synapses (Supplemental Fig. 5A-B) and the accumulations of syntaxin-3 with large vesicular structures (Supplemental Fig. 5B), we also observed perinuclear inclusions of syntaxin-3 in the ONL of Munc18-1 cKO cells (Supplemental Fig. 5C).

**Figure 5.**
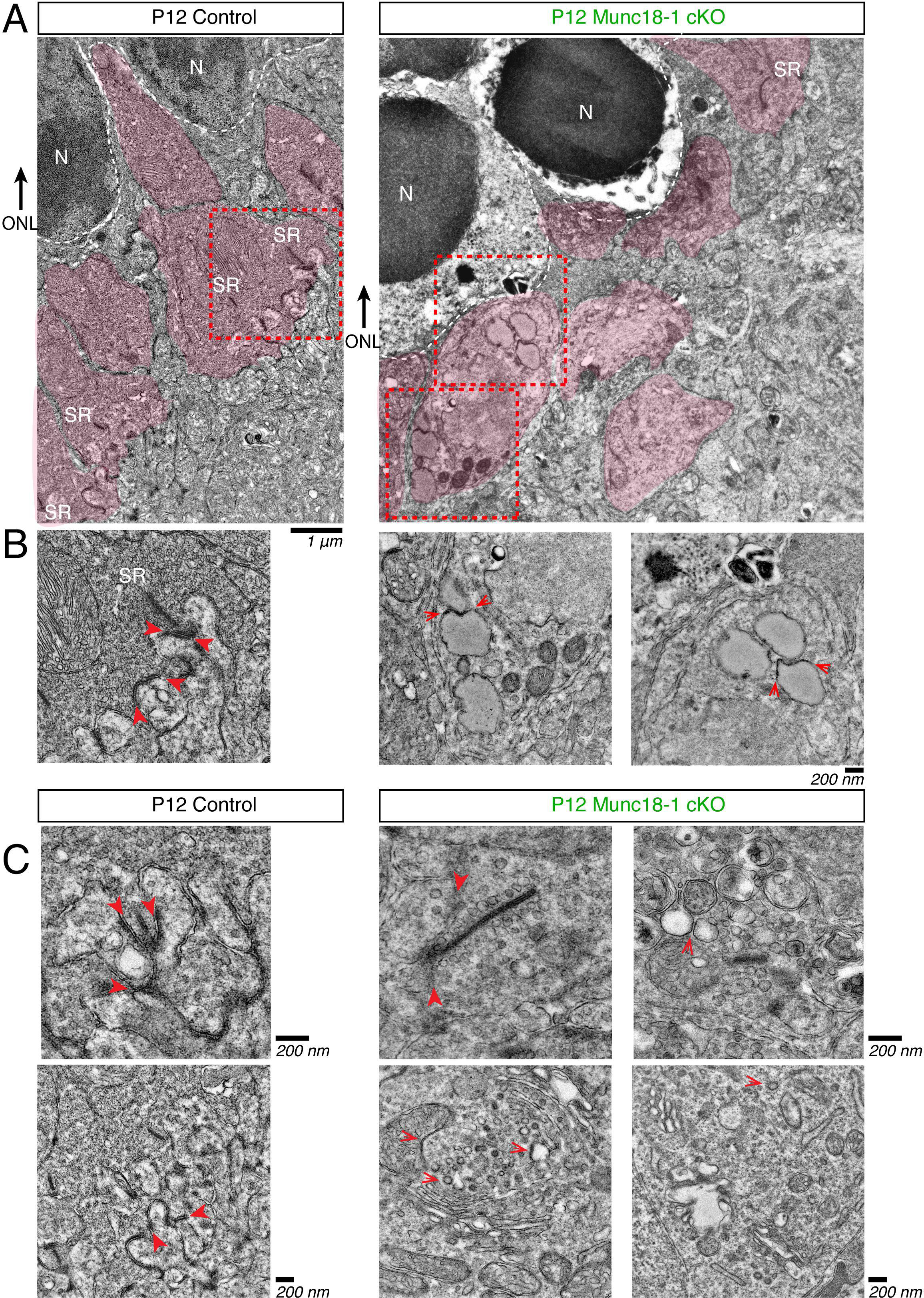
Syntaxin-3 is trapped in organelle membranes in Munc18-1 deficient photoreceptors. (A) Immunogold electron microscopy of control and Munc18-1 cKO retina. Individual photoreceptor cell terminals are pseudocolored in red. Syntaxin-3 is transported to the synaptic membrane in control photoreceptor terminals. In Munc18-1 deficient photoreceptor, synaptic ribbons (SR) are not found in close proximity to dark syntaxin-3 immunogold signal, rather syntaxin-3 signal is aggregated in non-synaptic membranes. (B) Magnified regions from (A). Control photoreceptors have developing synapses where the membrane is marked by syntaxin-3 immunogold labelling (red arrowhead). In Munc18-1 cKO terminals, syntaxin-3 signal is present on the membranes of lipidic structures (red arrows), mitochondria, and endoplasmic reticulum-like structures. (C) Further examples of syntaxin-3 presence on photoreceptor terminal synaptic membrane in control retina (red arrowhead). In Munc18-1 cKO, syntaxin-3 signal can also be seen on vesicles, mitochondria, vacuolated structures, and various organelles (red arrows).

We additionally did transmission electron microscopy to confirm changes at the ultrastructural level in P12 Munc18-1 control and cKO retinas. We observed skinny photoreceptor outer segments with neatly stacked membranous discs in the control sample, however, photoreceptor outer segments were often found to be swollen and vacuolated in the Munc18-1 cKO retina (Fig. 6A). These knockout outer segments also contained mitochondria and Golgi/endoplasmic reticulum appearing structures which are typically features of the photoreceptor inner segment (Fig. 6A). At the synaptic terminals, in the control, the synaptic boundary could be observed with developing photoreceptor synapses containing one synaptic ribbon per developing tripartite synapse. However, in Munc18-1 deficient photoreceptor cells, the synaptic boundary included vacuolated interruptions and dying photoreceptor nuclei could be observed (Fig. 6B). Furthermore, key synaptic features were altered – rather than a single synaptic ribbon per synapse, instances where multiple adjacent synaptic ribbons were present, some synaptic ribbon were found perinuclearly, and less synaptic ribbons were observed as a whole, confirming the presence of synaptic deficits in Munc18-1 cKO mice.

**Figure 6.**
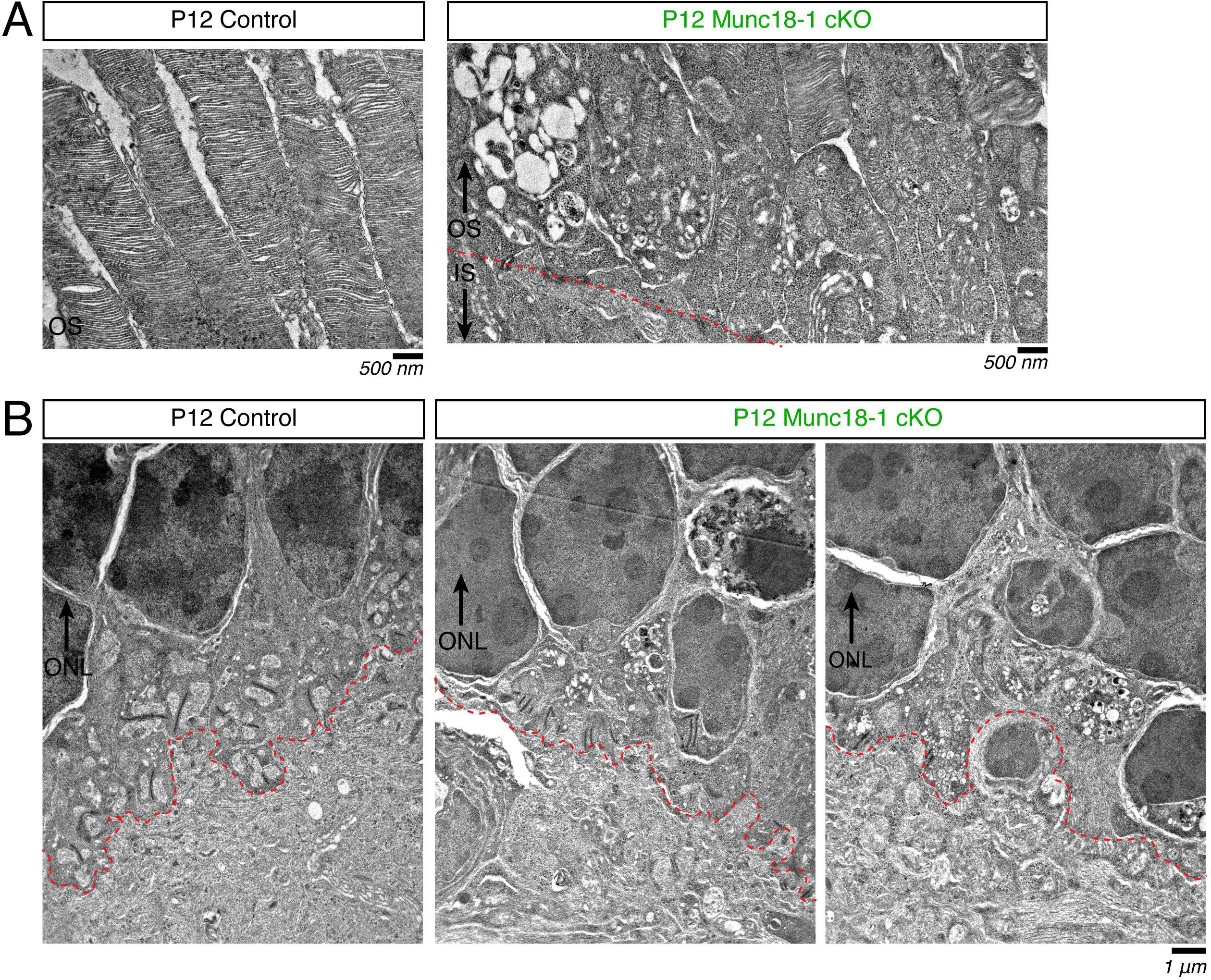
Outer segment and synaptic changes seen at ultrastructure level in Munc18-1 cKO retina. (A) Transmission electron micrographs of photoreceptor outer segments from P12 control and Munc18-1 cKO retina. Munc18-1 cKO outer segments are much more swollen and vacuolated compared to the skinny ciliary outer segments observed in controls. (B) Transmission electron micrographs of photoreceptor synaptic terminals. Developing photoreceptor terminals in control sections have many synaptic ribbons and clearly defined postsynaptic connections. Synaptic region (lined in red) has much more vacuolations and disturbances in Munc18-1 cKO retina. A loss of nuclear integrity is also observed in Munc18-1 cKO photoreceptors.

## Discussion

Here, we take advantage of the well-defined organization of the elegant retina system to investigate the complexity of Munc18-1 interactions with SNARE proteins in photoreceptor cells. To do so, we used CRX-cre to specifically remove Munc18-1 from photoreceptor cells and then investigated changes in function, morphology, and protein localization. Removal of Munc18-1 led to decreases in retina thickness largely between postnatal day 14 – 22 and diminished electroretinogram responses (Fig. 1). Notably, this 2-3-week window before Munc18-1 cKO retinas degenerated gave us the opportunity to investigate the interactions of Munc18-1 and syntaxin-3, and the targeting of SNARE proteins to photoreceptor presynaptic terminals the absence of Munc18-1. In Munc18-1 deficient photoreceptors, we found changes to syntaxin-3 localization – syntaxin-3 remained present in photoreceptor outer segments and was targeted towards synaptic terminals, but plasmalemmal located syntaxin-3 was significantly decreased with additional perinuclear occlusions of syntaxin-3 (Fig. 2). Despite syntaxin-3 being targeted towards synapses, we surprisingly found a differential location in synaptic syntaxin-3 relatie to its cognate t-SNARE partner SNAP-25 at photoreceptor synapses (Fig. 3). However, this effect was specific to syntaxin-3 as SNAP-25 and synaptobrevin-2 localization remained unchanged, and therefore, not an artefact of Munc18-1 removal interfering with synaptic proteins in general (Fig. 4). Using immuno-EM, we were able to confirm syntaxin-3 presence in photoreceptor synaptic membranes of the control retina, but in the Munc18-1 cKO retina, syntaxin-3 localization was observed in organelle membranes, vesicles, and vacuoles (Fig. 5). Using transmission electron microscopy, we were also able to observe the early degeneration of photoreceptor outer segments and synaptic terminals (Fig. 6).

Our results indicate an essential role of Munc18-1 in photoreceptor cell survival and synaptic function. Recently, syntaxin-3 and SNAP-25 were determined to be the obligate t-SNARE proteins photoreceptor cells use in synaptic transmission and outer segment protein trafficking (Kakakhel et al., 2020; Janecke et al., 2021; Huang et al., 2024). Thus, we conclude the photoreceptor ribbon synapses utilize not only SNARE protein dependent but also sec/Munc18 protein dependent mechanisms for synaptic functions.

While syntaxin-3 and SNAP-25 are both enriched in control photoreceptor synaptic terminals, the localization of syntaxin-3 and SNAP-25 is strikingly segregated in the Munc18-1 deficient nerve terminals. This unexpected result suggests that Munc18-1 is essential for ensuring syntaxin-3 is transported to photoreceptor terminals and for its co-localization with SNAP-25 (Fig. 7). Can this finding extend to more conventional neurons and synapses in the brain? In conventional synapses, rather than syntaxin-3, syntaxin-1A and syntaxin-1B, are the key isoforms for synaptic vesicle exocytosis. Our results that syntaxin-3 is transported to terminals of Munc18-1 deficient photoreceptors seem to be consistent with previous results that syntaxin-1 was correctly targeted to synapses in embryonic day 18 (E18) Munc18-1 null brain slices (Toonen et al., 2006). Our finding of perinuclear aggregation of syntaxin-3 in the soma of photoreceptors is also consistent with cytoplasmic accumulation of syntaxin-1 in the soma of neurons that are differentiated from patient-derived Munc18-1 heterozygous nonsense mutant iPSCs (Yamashita et al., 2016). Thus, we speculate that syntaxin-1 is also segregated from SNAP-25 within Munc18-1 deficient conventional synapses.

**Figure 7.**
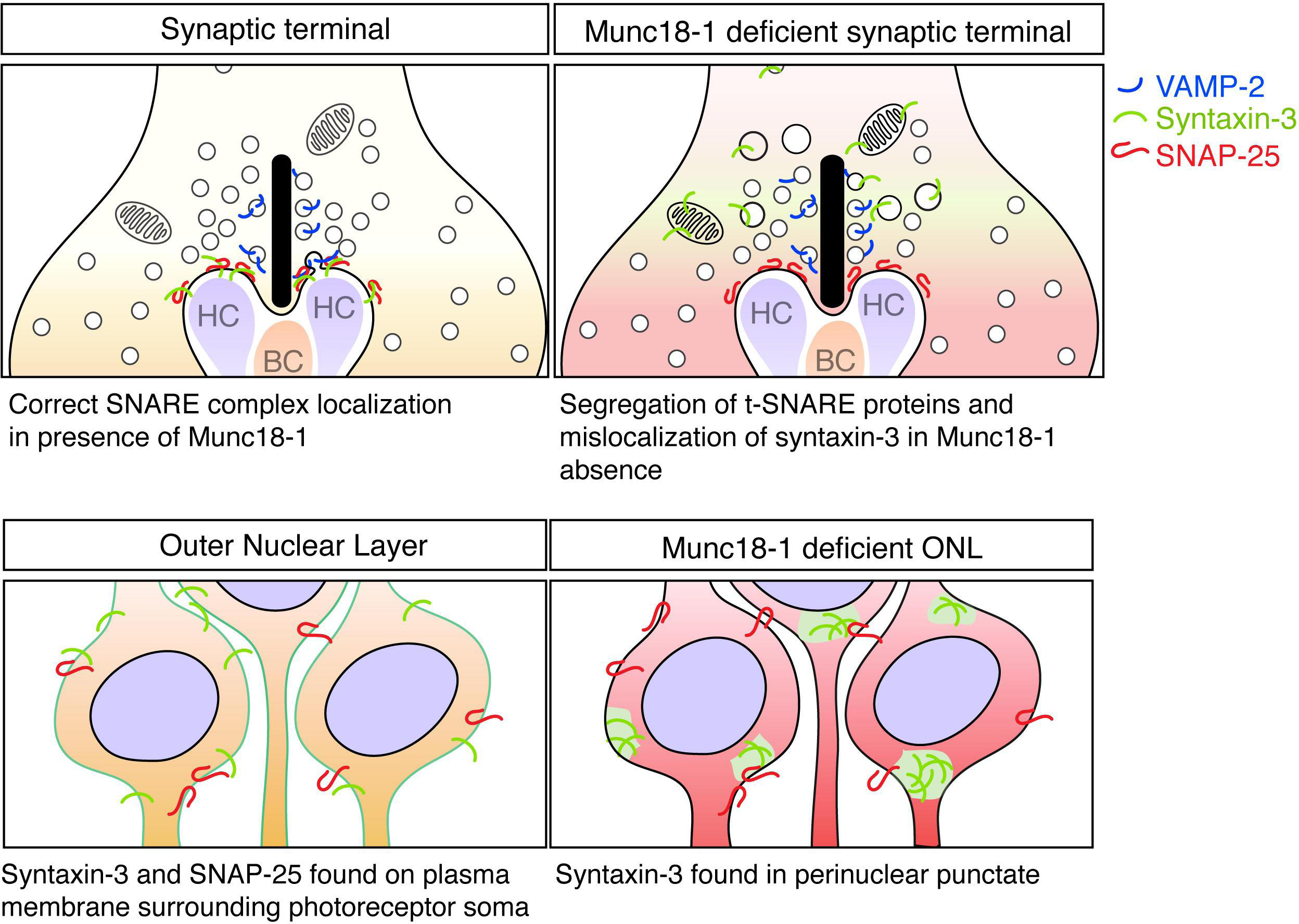
Segregation of t-SNARE proteins and mislocalization of syntaxin-3 in Munc18-1 absence.

Although Munc18-1 has been recognized as a regulator of neurotransmitter exocytosis, its role is considered to extend past regulation for exocytosis. For instance, Munc18-1 removal results in the complete absence of both spontaneous and evoked release (Verhage et al., 2000), which is more severe than the phenotype of the knockout of synaptobrevin-2 or SNAP-25 (Schoch et al., 2001; Deák et al., 2004; Washbourne et al., 2002; Bronk et al., 2007). While spontaneous release is still present in synaptobrevin or SNAP-25 knockouts, there is no spontaneous release in Munc18-1 knockout. The severe phenotype of Munc18-1 absence is more comparable to the double knockout of syntaxin-1A and 1B, which is embryonic lethal (Vardar et al., 2016; Kofuji et al., 2014). The essentiality of Munc18-1 homologues is also demonstrated in lower organisms such as *Drosophila* and *C. elegans*. Rop, the *Drosophila* homologue of Munc18, is essential for the life of *Drosophila* and its null mutation results in embryonic lethality (Harrison et al., 1994). Meanwhile, the *C. elegans* homologue of Munc18 is known as UNC-18, and while unc-18 null *C. elegans* are viable, recent results suggest that this viability is supported by the presence of another isoform of Munc18, UNCP-18, and a double mutation of *unc-18* and *uncp-18* results in embryonic lethality (Boeglin et al., 2023). Such severe phenotypes observed in Munc18 mutants do not seem to be explained by mere regulatory roles of Munc18-1 in the SNARE complex (Han et al., 2010). Previous studies emphasise the regulatory role of Munc18-1 after t-SNARE complex assembly or in the acceleration of tertiary SNARE complex assembly (Baker et al., 2015; Park et al., 2017; Sitarska et al., 2017; Parisotto et al., 2014), and such regulatory role of Munc18-1 seem to be mediated by its domain 3 (Han et al., 2013; Munch et al., 2016; Park et al., 2017; Meijer et al., 2012). However, our results suggest that Munc18-1 is essential for syntaxin to be localized to the same membrane as SNAP-25, namely at the presynaptic plasma membrane.

The remaining question is why in the absence of Munc18-1, syntaxin-3 is trafficked to photoreceptor terminals, but remains absent from the plasma membrane. It has been suggested that syntaxin is notorious in interacting with numerous proteins and is a very promiscuous protein (Wu et al., 1999; Lewis et al., 2001; Wendler and Tooze, 2001); syntaxin-1 also has a known tendency to oligomerize and cluster, and this property is likely shared by syntaxin-3. Thus, without the protection by Munc18-1, it may aggregate with itself or bind with other synaptic proteins, hampering its arrival to the terminal plasma membrane, and instead becoming trapped in various organelle membrane structures. It has also been previously proposed that that syntaxin-3 has a lower affinity for SNAP-25 (Liu et al., 2014; Nishad et al., 2023), which may indicate syntaxin-3 may mostly exist in the closed conformation, though our results may suggest that the lowered affinity of syntaxin-3 for SNAP-25 may only be partially attributed to Munc18-1 stabilizing a closed conformation of syntaxin-3. Alternativity, Munc18-1 may provide the mechanisms to transfer syntaxin away from intramembrane structures to the terminal plasma membrane. In any case, our finding suggest that Munc18-1 is critical for the proper localization of syntaxin-3 to the presynaptic membrane of photoreceptor terminals, and syntaxin-3 and SNAP-25 co-localization is lost in the absence of Munc18-1.

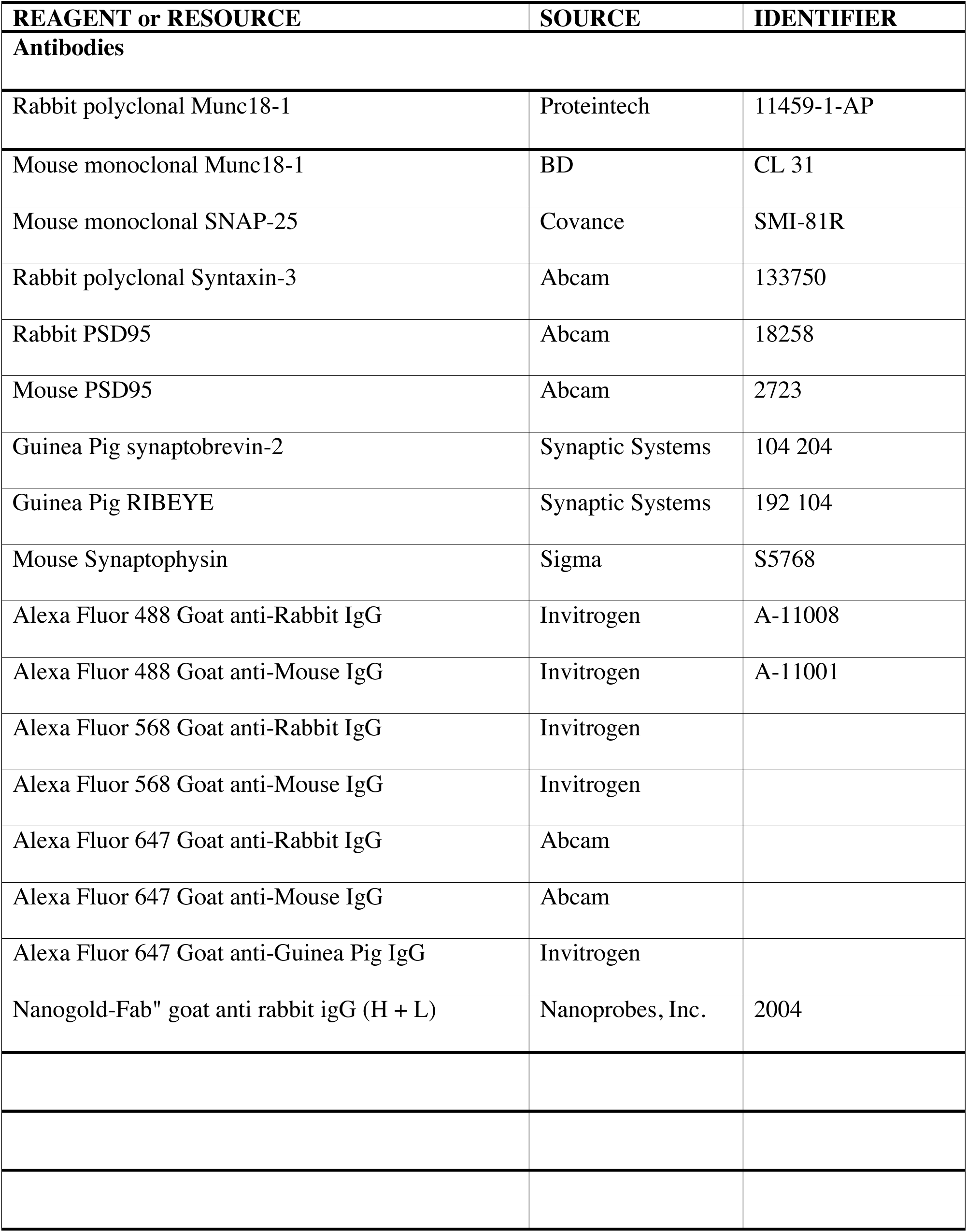
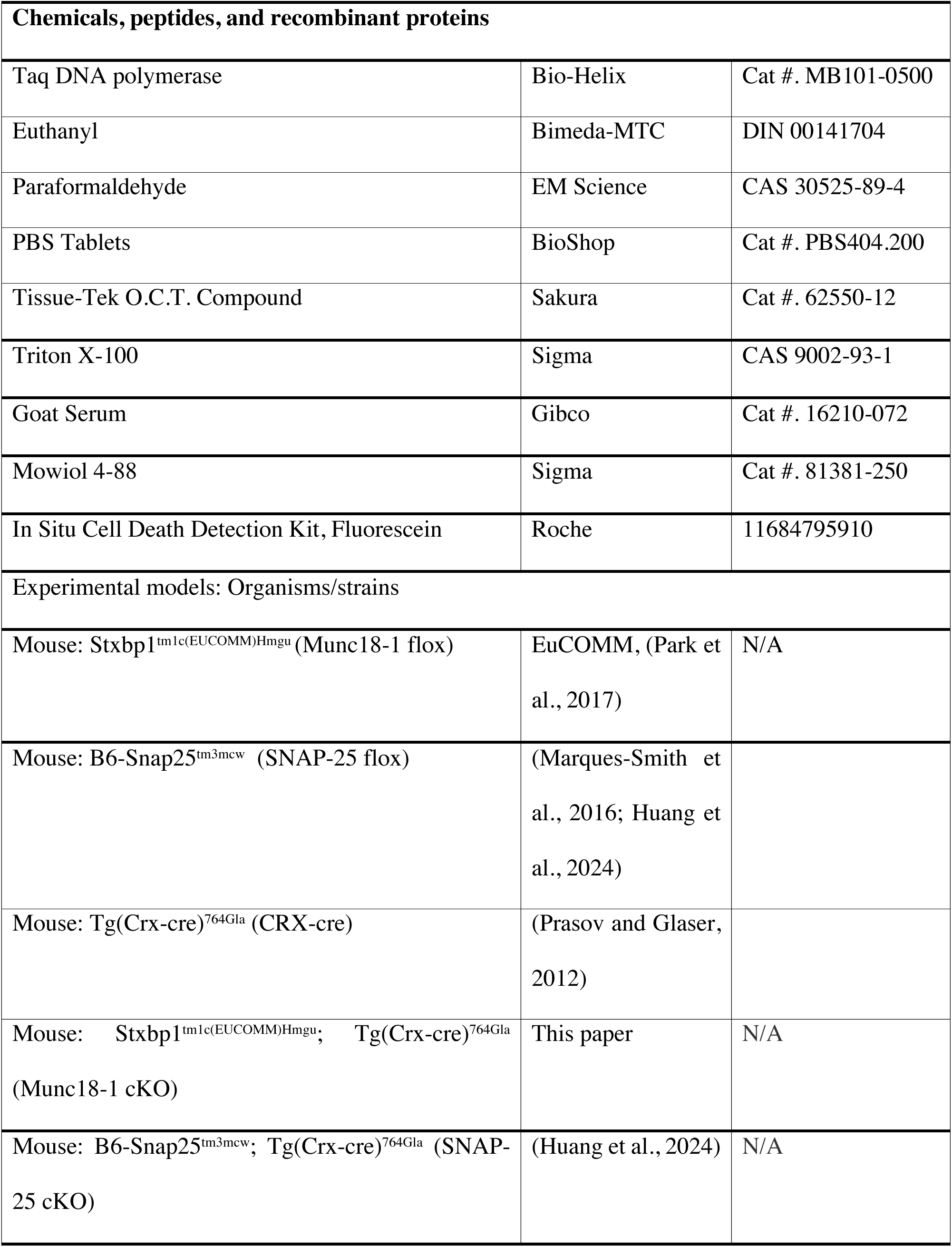
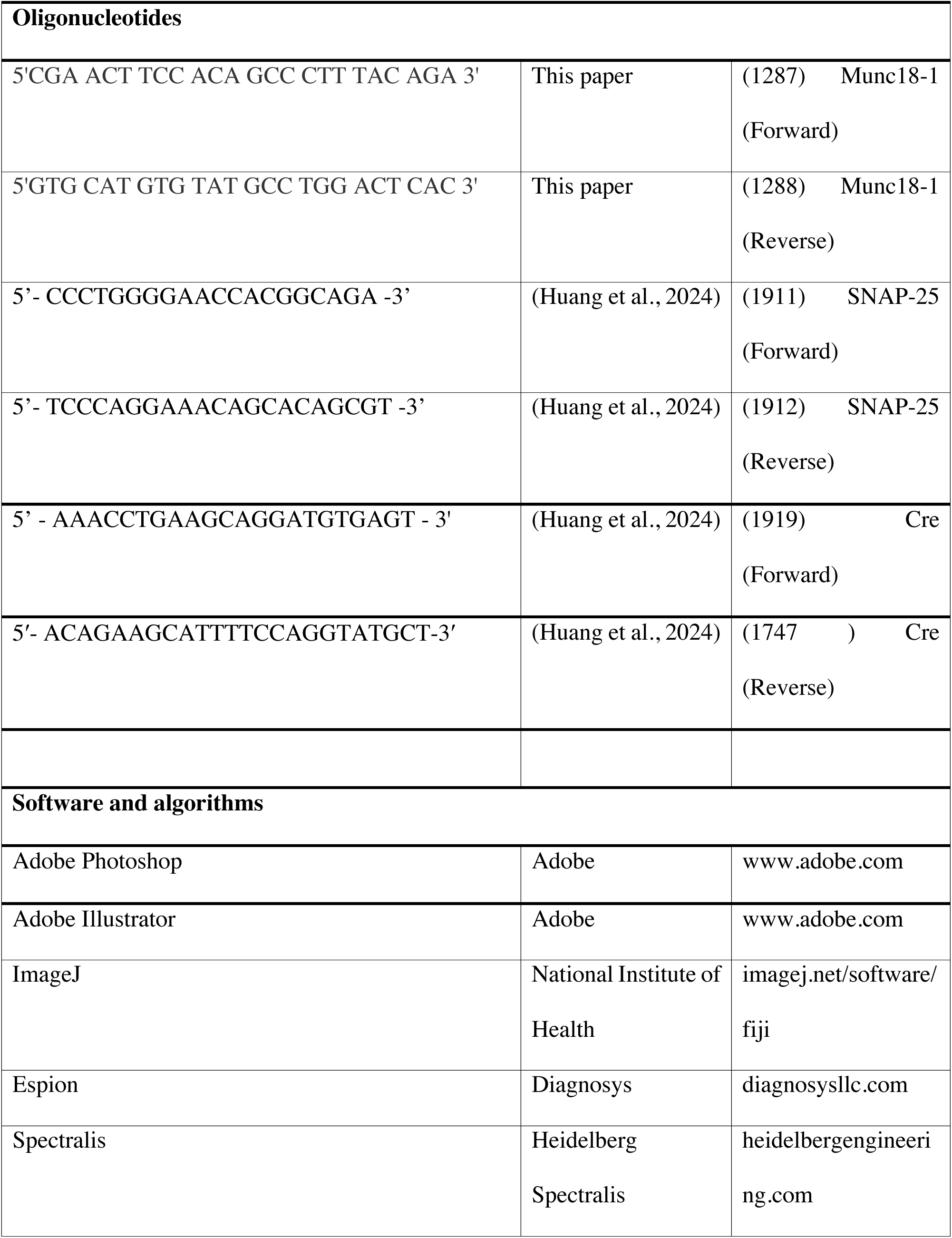
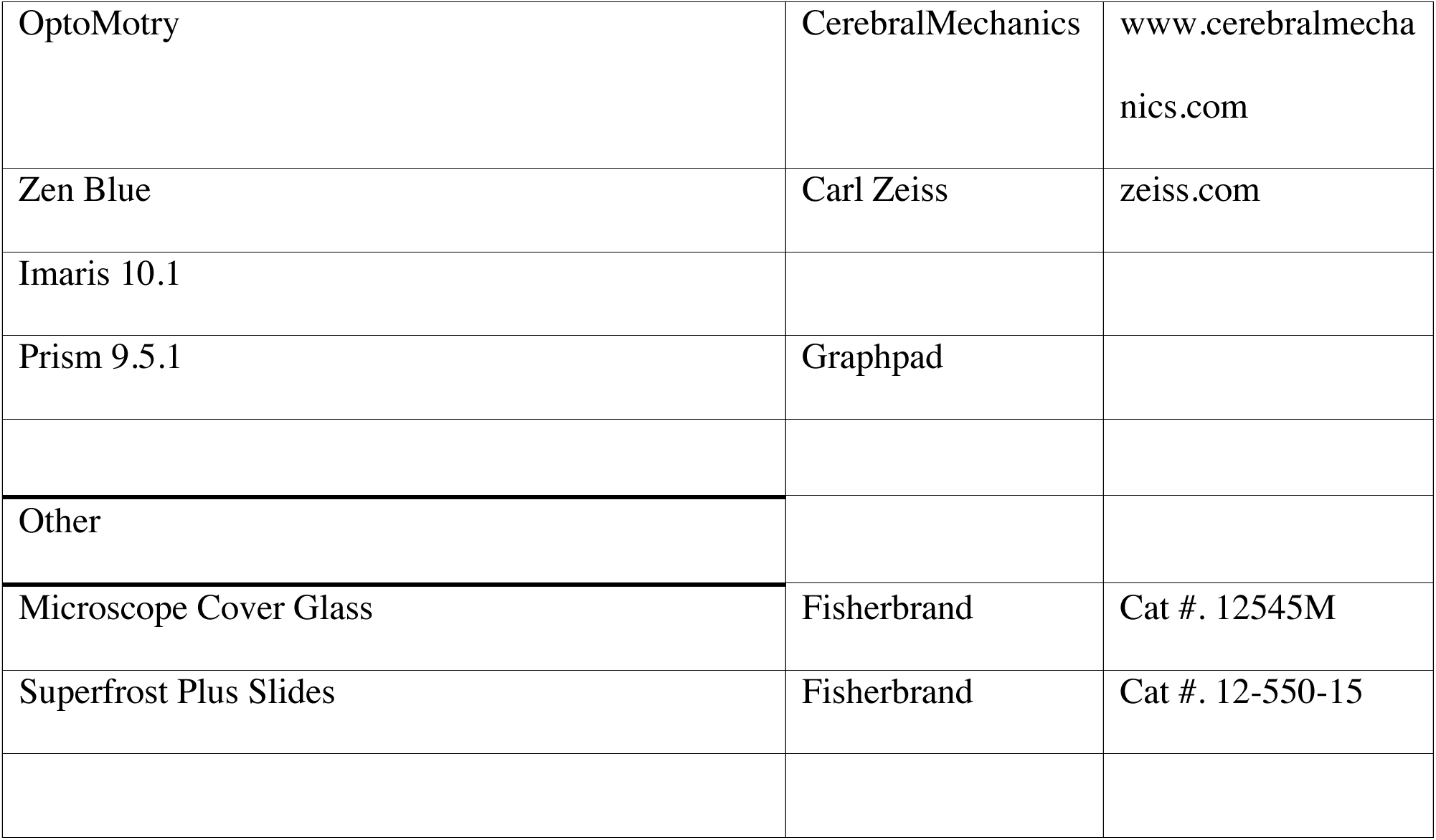

## DATA AVAILABILITY

### Lead contact

Further information and requests for resources and reagents should be directed to and will be fulfilled by the lead contact, Shuzo Sugita.

## Materials availability

All reagents and mouse lines generated by this study are available from the lead contact

## METHODS

### Experimental model and subject details

Munc18-1 and SNAP-25 flox mice (Hoerder-Suabedissen et al., 2019; Marques-Smith et al., 2016; Huang et al., 2024; Park et al., 2017) of both sexes between postnatal day 9 and up to 2 months of age were used in this study. SNAP-25 flox and conditional mice were generated as previously described (Huang et al., 2024), briefly, For the flox mice, loxP sites were introduced to surround exon 5 of SNAP-25 in SNAP-25 flox mice and SNAP-25 flox mice were crossed with CRX-cre mice (https://www.informatics.jax.org/allele/MGI:5433291) to obtain SNAP-25 cKO mice (Huang et al., 2024; Prasov and Glaser, 2012). For Munc18-1 flox mice, loxP sites were introduced surrounding exon 7, and Munc18-1 conditional knockout mice were generated by crossing Munc18-1 flox mice with CRX-cre mice. Genomic DNA was obtained from tail clippings using alkaline lysing methods and animals were genotyped using PCR (refer to table for primers). Tail clippings were lysed with 50 mM NaOH and incubated at 96 °C for 1 hour with vigorous shaking. Littermate flox/flox mice without CRX-cre were used as controls for all experiments. All mice were maintained on a C57BL/6 background.

All mice were housed in a vivarium with a 12-hour light on/off cycle and temperatures were maintained between 22 – 23°C. We have complied with all relevant ethical regulations for animal use. All experiments detailed here were reviewed and approved by the animal care committee of the University Health Network in accordance with the Canadian Guidelines for Animal Care.

### Preparation of retinal samples for immunohistochemistry

Mice were anaesthetized with an intraperitoneal injection of sodium pentobarbital (75 mg/kg, Bimeda-MTC) and subsequently euthanized. Eyes were placed in 4% PFA following enucleation.

Using a 32G syringe, a hole was poked in the cornea and to aid in the fixation process and eyes were postfixed for 2 hours at room temperature in 4% PFA. Following, three PBS washes, the eyes were placed in 30% sucrose overnight at 4°C for dehydration. The next day, eyes were mounted in OCT and stored at -80°C until sectioning. 14 μm sections were obtained using a Leica CM3050S cryostat. Sections were dried and stored at -80°C until use.

### Preparation of retina samples for transmission electron microscopy

Mice were euthanized as previously described and transcardially perfused with 5 – 10 mls of PBS followed by EM fixative. Eyes were enucleated and placed into EM fixative. The front of the eye was removed but the retina and all extra scleral tissue was left intact. An approximately 4 mm x 4 mm square was cut from the mid-periphery of the eye for further processing. Further sample processing for transmission electron microscopy were performed by the Nanoscale Biomedical Imaging Facility, The Hospital for Sick Children, Toronto, Canada. Transmission electron micrographs were taken on a Hitachi HT7800 transmission electron microscope.

### Preparation of retina samples for pre-embedding immuno-electron microscopy

Mice were euthanized as previously described and transcardially perfused with 5 – 10 mls of PBS followed by 5 – 10 mls of fixative solution (4% PFA + 0.1% glutaraldehyde) for 3 – 5 mins. Eyes were enucleated and placed into fixative. A 32G syringe was used to poke a hole in the cornea and an opening was cut in the front of the eye to allow fixative to enter the eye cup. The lens was removed and the eyecup was fixed for 1 hour. The eyecup was then washed three times using PBS for 5 minutes each. The eyecup was embedded in low-gelling temperature agarose and 150 μm sections was cut using a vibratome and PBS as a buffer. Retina sections were collected into a well plate with PBS until further use.

Sections were permeabilized and blocked using 10% goat serum + 0.05 triton X-100 in PBS on a shaker for 30 minutes. After 30 minutes, the blocking solution was removed and the primary antibody solution (1:1000 syntaxin-3 antibody in PBS + 1% goat serum + 0.005% triton X-100 was added to the sections. Sections were incubated in primary antibody solution with gentle shaking at 4°C for 4 days.

After primary incubation, sections were washed 3 times in PBS + 1% goat serum for 5 minutes each. Secondary antibody solution (1:100 anti-rabbit 1.4 nm nanogold in PBS + 1% goat serum + 0.005% triton X-100) was applied and sections were incubated for 2 hours. Sections were then washed twice with PBS + 1% goat serum, and twice more with PBS. Finally, sections were placed in fixative and stored until further processing. Further sample processing for transmission electron microscopy were performed by the Nanoscale Biomedical Imaging Facility, The Hospital for Sick Children, Toronto, Canada. Transmission electron micrographs were taken on a Hitachi HT7800 transmission electron microscope.

### Immunohistochemistry

Retina sections were washed three times for 3 minutes each in PBS followed by three 5-minute washes in PBS with 0.1% triton X-100 (PBS-Tx). Sections were then blocked with 10% goat serum in PBS-Tx for 1 hour. Primary antibody was diluted 1:500 in blocking solution and left overnight at 4°C in a humidified chamber. The following day, five by 5-minute PBS-Tx washes were done, and the secondary antibody solution (1:1000 antibody in with DAPI+PBS-Tx) was applied to sections for 1 hour at room temperature and covered from light. Further antibody detail and information are provided in the attached resource table. Sections were then washed five times using 5-minute PBS-Tx washes, while covered from light and with shaking before sections were placed in PBS and mounted in Mowiol.

### Immunofluorescence imaging

Sections were imaging using a Carl Zeiss confocal microscope (LSM780) and with a minimum averaging of 4x. Images were taken at 20x and 63x using 1.0 Airy Unit pinhole sizes. For 63x oil immersion images, Zeiss immersion oil 518F was used. A region 0.5 mm – 1.0 mm away from the optic nerve was selected for imaging and displayed in all representative immunofluorescence images.

### Image processing and quantification

For profile analysis, line intensity histograms were generated in ImageJ. For each image, three regions were selected and within each region, three 5 μm wide line profiles were generated. The intensity of the syntaxin-3 and SNAP-25 channels were generated along each pixel of the line. The generated profiles were then aligned to the peak intensity of syntaxin-3. The three 5 μm wide line profiles within each region was then averaged and the resulting separated signal intensities for each dye were plotted as a function of distance.

For volume analysis, images were rendered in Imaris 10 (Bitplane, Zurich). Using the surfaces function, a surface of a 500 px wide, 100 px tall region of the outer plexiform layer was generated for both syntaxin-3 and SNAP-25 channels.

### Optical coherence tomography

Spectral-domain optical coherence tomography scans were performed on mice of 3 weeks of age using the Heidelberg Spectralis animal imaging system. Mice were anesthetized using Avertin (125-250 mg/kg) and had their eyes dilated using Mydriacyl. The left eye of each mouse was used for all scans and all thickness measurements were taken 1.0 – 1.5 mm away from the optic nerve head, automated segmentation provided by the Heidelberg Spectralis animal imaging software and the experimenter manually fixed erroneous segmentation. Each round of scanning was performed in under 10 minutes and mice were let to recover on a heated pad.

### Electroretinograms

Electroretinogram recordings were performed on P22 mice. Light stimulation was through the Diagnosys Espion electroretinogram device with a ColorDome Ganzfeld stimulator and Espion software. Before recordings, mice were dark adapted overnight and all recordings were done under red illumination. Mice were anesthetized using Avertin (125-250 mg/kg). Mydriacyl (Alcon) eye drops were used to dilate the eyes, and topical Alcaine (Alcon) was also applied. Body temperature was maintained at 38°C throughout examination using the Diagnosis device. Recordings were taken from both eyes using 2 mm gold loop electrodes placed on the cornea of the eyes. Reference electrode was placed between the eyes subcutaneously and a ground electrode in the tail. The cornea was kept moist with a thin layer of methylcellulose solution. Light stimulation consisted of brief pulses of white light (6500K). Scotopic recordings were done first on the dark-adapted mouse. 10 recordings per stimulus intensity were averaged, and the a- and b wave amplitudes were manually determined by the experimenter. Scotopic stimuli conditions are as follow: 0.00025 cd/s/m^2^, 0.0025 cd/s/m^2^, 0.025 cd/s/m^2^, 0.25 cd/s/m^2^, 2.5 cd/s/m^2^, 5 cd/s/m^2^, and 10 cd/s/m^2^. Mice were then light adapted with 30 cd/s/m2 background illumination for 10 minutes. Photopic tests were done in the presence of 30 cd/s/m2 background illumination. 10 recordings per flash intensity were averaged, and the amplitude of the a- and b waves was manually determined by the experimenter again. The following photopic stimulus intensities were used: 0.15 cd/s/m^2^, 1.25 cd/s/m^2^, 5 cd/s/m^2^, 10 cd/s/m^2^, 25 cd/s/m^2^, 50 cd/s/m^2^, and 75 cd/s/m^2^.

### Optomotry testing

Optomotry testing was performed on Munc18-1 control and cKO mice of 2 months of age. Quantification of visual acuity was performed using the OptoMotry system from CerebralMechanics. Vertical sine wave gratings were presented by a chamber composed of four monitors to create a virtual cylinder in the visual field of the mouse. A Zeiss video camera was used to image the mouse from above. The mouse’s head was manually tracked by a software and the vertical sine wave gratings were then rotated around the mouse at 12 degrees/second using the randomized protocol design and the simple staircase psychophysical method within the software. The tester was blinded to spatial frequencies being tested during the procedure.

### Statistics and Reproducibility

All statistical analyses were done in Prism 9.5.1. Independent t-tests were used for two-group experiments with a p-value < 0.05 as the threshold for statistical significance. For all quantified tests (OCT, ERG, OKT), minimum triplicates were used at every time point. No data were excluded from analysis. Reproducibility was confirmed using triplicates. Two-way analysis of variance (ANOVA) was used for comparisons of multiple groups, with a significance level of 0.05. All error bars represent SEM.

## Acknowledgements

We thank Josep Rizo (UT Southwestern) for critically reading an earlier version of our manuscript. This work was supported by the Natural Sciences and Engineering Research Council of Canada (RGPIN 2020 07139) and the Canadian Institutes of Health Research (CIHR PJT 165917). M.H. and C.H.C. are supported by the Vision Science Research Program scholarship, Canada Graduate Scholarship, and the Ontario Graduate Scholarship. The authors wish to thank the Nanoscale Biomedical Imaging Facility, The Hospital for Sick Children, Toronto, Canada for assistance with transmission electron micrographs and immuno-EM.

