## Supplemental Files for "Segregated localization of target-SNARE proteins within presynaptic terminals of Munc18-1 deficient photoreceptors"

**Supplemental Figures.**

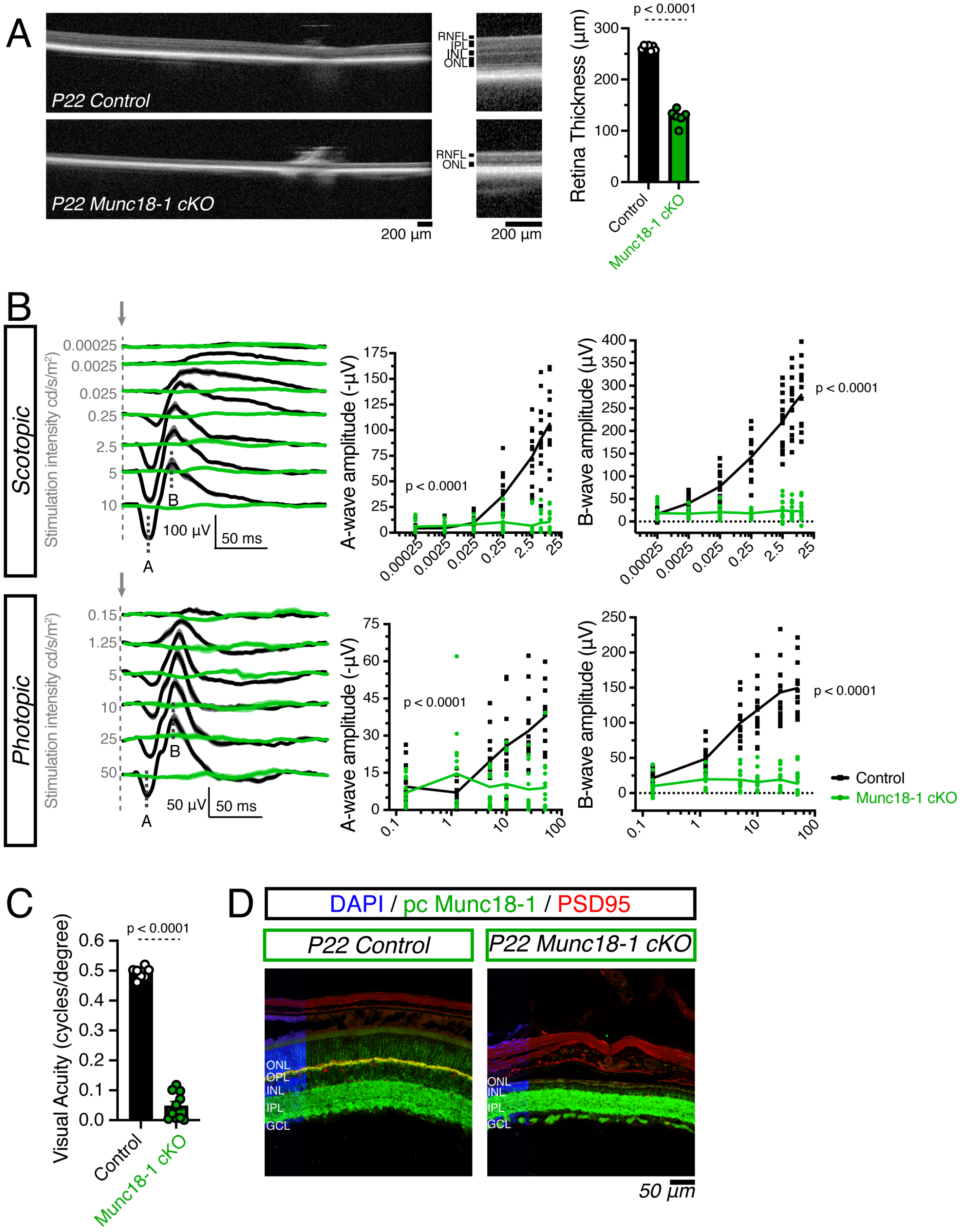

**Supplemental Figure 1. Phenotype of Munc18-1 cKO mice.**

(A) Optical coherence tomography live imaging of P22 control and P22 Munc18-1 cKO retinas with quantifications. A significant decrease in retinal thickness is observed in Munc18-1 cKO retinas; $t_{(11)} = 22.47$; $p < 0.0001$. Scale bar = 200 μm

(B) Averaged scotopic (dark-adapted) and photopic (light-adapted) electroretinogram traces across 7 light intensities in P22 control and P22 Munc18-1 cKO mice. Arrow indicate stimulation onset, A indicates a-wave, B indicates b-wave. Munc18-1 cKO mice exhibit no responses to light stimulation at any light intensity.

Quantifications of amplitudes. Scotopic A-wave amplitudes: two-way ANOVA; $F(1, 26) = 69.03$; $p < 0.0001$; scotopic B-wave amplitudes: two-way ANOVA; $F(1, 26) = 139.3$; $p < 0.0001$. Quantification of photopic A-wave amplitudes: two-way ANOVA; $F(1, 26) = 21.15$; $p < 0.0001$; photopic B-wave amplitudes: two-way ANOVA; $F(1, 26) = 125.5$; $p < 0.0001$. Calibration bars represent 50 ms (horizontal). Vertical calibration bars are 100 μV (scotopic) and 50 μV (photopic).

(C) Visual acuity assessed by measuring optokinetic tracking response using CerebralMechanics OptoMotry system. Severe and significant decrease to visual acuity is observed in Munc18-1 cKO mice compared to control following Munc18-1 removal from photoreceptors, $t_{(15)} = 24.07$; $p < 0.0001$.

(D) Antibody staining for Munc18-1 (green) using rabbit polyclonal antibody against PSD95 (red) for presynaptic terminals in P22 control and P22 Munc18-1 retina. P22 control retina shows strong Munc18-1 signal throughout while Munc18-1 cKO retina maintains Munc18-1 through the inner nuclear layer, inner plexiform layer and ganglion cell layer. Munc18-1 cKO outer nuclear layer is severely degenerated and virtually no photoreceptors remain by P22.

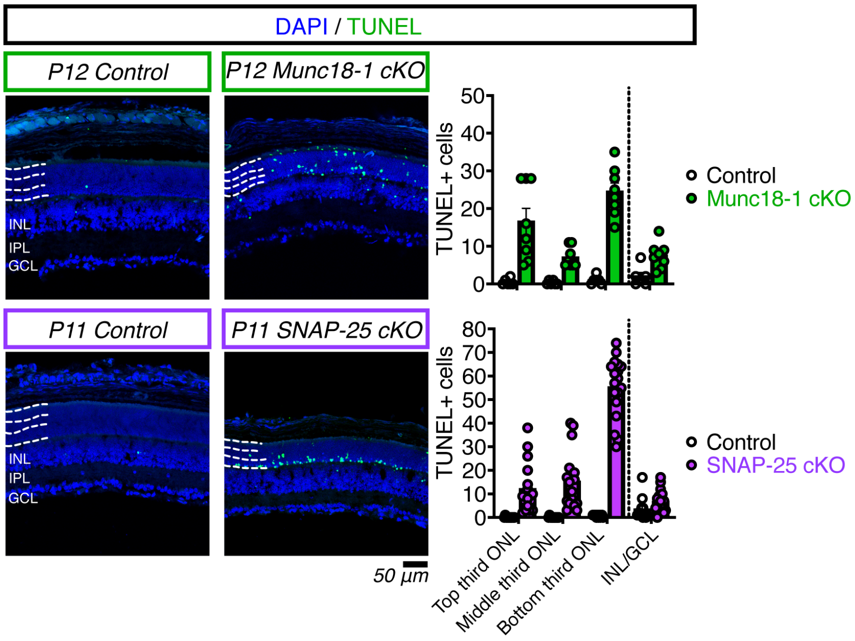


**Supplemental Figure 2.**

TUNEL staining (green) in control, Munc18-1 cKO (green) and SNAP-25 cKO (purple) retina. When dividing the outer nuclear layer (ONL) into thirds, SNAP-25 cKO mice (purple) show a lower bias for TUNEL signal, suggesting photoreceptors are dying from the lowermost strata. Munc18-1 cKO have a different TUNEL pattern where there is increased TUNEL signal in the top third and bottom third of the ONL.

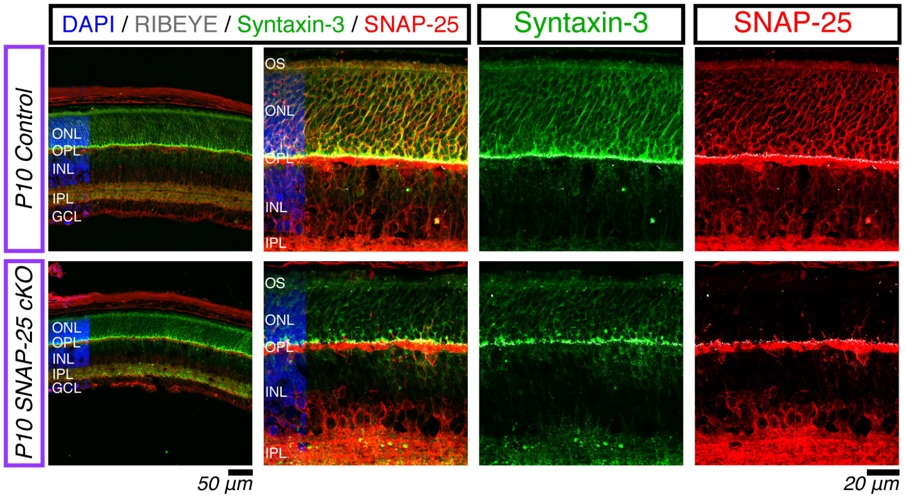


**Supplemental Figure 3.**

Syntaxin-3 (green) and SNAP-25 (red) immunostaining in P10 control and SNAP-25 cK retinas. Despite the severe synaptic degenerations observed in SNAP-25 cKO animals, syntaxin-3 is nevertheless correctly located in the outer segments, surrounding photoreceptor nuclei, and in the synapse.

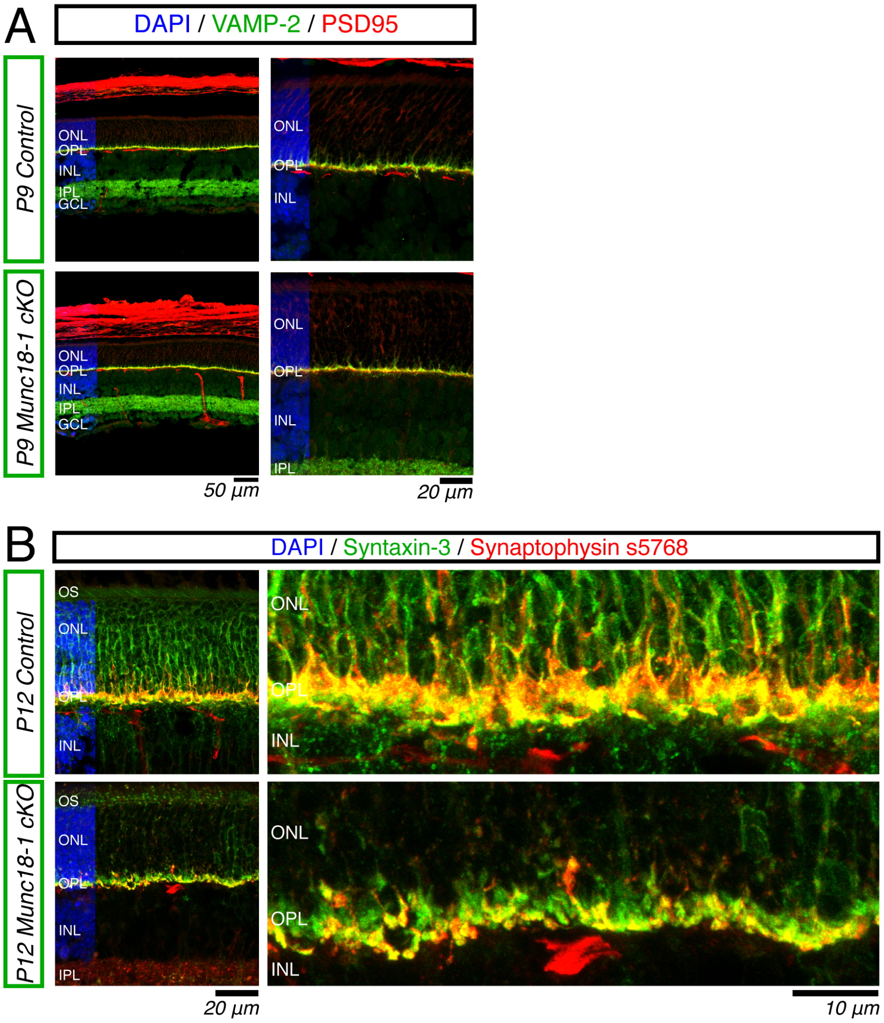


**Supplemental Figure 4.**

Synaptobrevin-2 (green) and PSD95 (red) staining in P9 control and Munc18-1 cKO retina. Synaptobrevin-2 colocalizes in presynaptic photoreceptors as identified by colocalization between synaptobrevin-2 and PSD95. Synaptobrevin-2 remains properly colocalized with PSD95 in Munc18-1 cKO retinas.

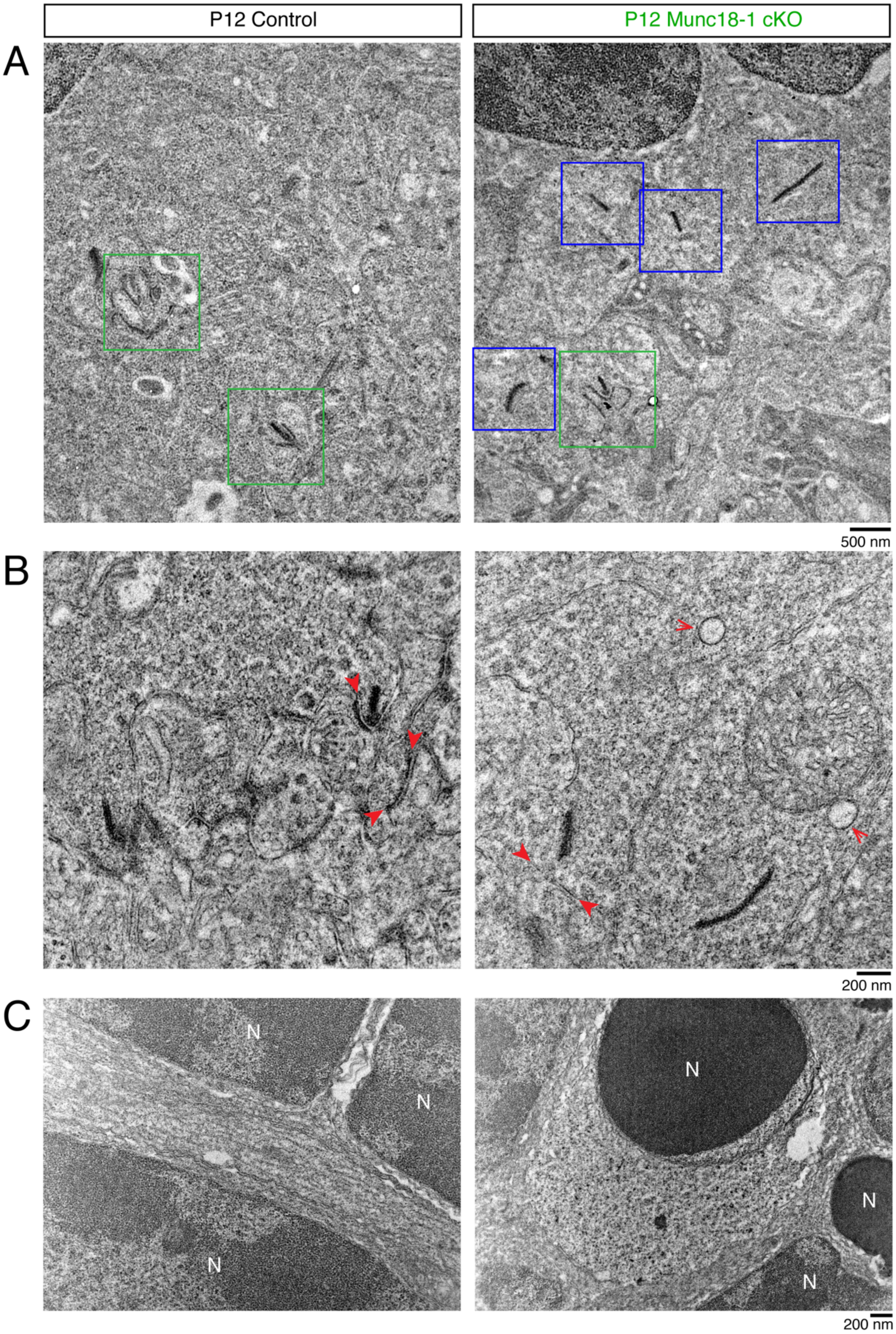

P12 Control
P12 Munc18-1 cKO
A
B
C
500 nm
200 nm
N
N
N
N
N
N
200 nm

**Supplemental Figure 5.**

Immuno-electron micrograph of P12 control and Munc18-1 cKO retina without uranyl acetate and lead citrate contrasting.

(A) Synaptic regions in P12 control and Munc18-1 cKO retina sections. Photoreceptor synaptic membranes adjacent to synaptic ribbons are dark and spotted with syntaxin-3 immunogold labelling (green box). In Munc18-1 cKO photoreceptors, one synaptic terminal is labelled by syntaxin-3 immunogold (green box), likely representing a cell where cre-recombination was less effective. Other four synaptic membranes adjacent to synaptic ribbons (blue boxes) all lack syntaxin-3 immunolabelling.

(B) Magnified synaptic region of P12 control and Munc18-1 cKO retina. Control photoreceptor membranes near synaptic ribbons are tagged by syntaxin-3 (red arrowhead), while Munc18-1 cKO retina synaptic membranes near synaptic ribbons lack syntaxin-3 signal (red arrowhead), and syntaxin-3 signal is present in vesicular structures (red arrows).

(C) Magnified outer nuclear layer in control and Munc18-1 deficient photoreceptors. Control photoreceptor nuclei are surrounded by plasma membrane with uniform labelling. In Munc18-1 deficient photoreceptors, perinuclear occlusions of syntaxin-3 can be observed in addition to a change in nuclear integrity identified by chromatin condensation in figure panel.
